# Phytoremediation of 25+ Years Weathered Petroleum Contamination with Symbiotic Arbuscular Mycorrhizal Fungi and *Bacillus subtilis* ATCC 21332

**DOI:** 10.1101/2025.08.22.671748

**Authors:** Prama Roy

## Abstract

Petroleum hydrocarbon (PHC) contamination affects approximately 60% of Canadian contaminated sites, with the boreal ecozone facing acute exposure from industrial activities. This study evaluated the long-term effectiveness of plant-microbe enhanced phytoremediation of 25+ years weathered PHC-contaminated field soil over a 24-month period. Five native or naturalized plant species (three tufted grasses, a forb, and a tree) were tested across two contaminated soils and a control to compare remediation efficacy and plant health: 25,700 mg/kg total petroleum hydrocarbons (TPH) and a 1:1 contaminated and background site soil mixture (12,600 mg/kg TPH). Soil treatments included a Plants-Only Control, Arbuscular Mycorrhizal Fungi (AMF), plant-growth-promoting-rhizobacterium - *Bacillus subtilis*, and AMF+*B. subtilis*. Plant inoculation significantly improved remediation compared to natural attenuation (No Plants Control) in both PHC-contaminated soils. AMF-based treatments significantly remediated both soils: AMF in the 25,700 mg/kg TPH soil (74% mean TPH remediation) and AMF+*B. subtilis* in the 12,600 mg/kg TPH soil (96% remediation). In the 12,600 mg/kg TPH soil, all treatments successfully reduced PHC levels to below federal and provincial thresholds (≥95% removal) within 17 months. *Picea mariana* (black spruce) and *Andropogon gerardii* (big bluestem) above- and below-ground biomasses were not significantly impaired by the 25,700 and 12,600 mg/kg TPH soils, indicating their suitability for site remediation. *B. subtilis* significantly improved above-ground biomass and plant height and is therefore recommended for co-inoculation with AMF. These findings demonstrate that AMF-enhanced phytoremediation represents a highly effective, sustainable approach for long-term PHC remediation in Canadian boreal ecosystems, offering significant advantages over natural attenuation processes alone.

**Highlights:**

- AMF-based treatments achieved highest PHC removal in 25+ year weathered boreal soils
- Prairie grasses remediated 12,600 mg/kg TPH soil by >95% in AMF+*B. subtilis* treatment
- Enhanced *Picea mariana* (black spruce) biomass and moisture in 25,700 mg/kg TPH soil
- *Bacillus subtilis* significantly enhanced plant above-ground biomass and height
- AMF helped maximize PHC remediation, while *B. subtilis* promoted plant growth

## 1.0 Introduction

Petroleum hydrocarbon (PHC) contamination represents one of the most pervasive environmental challenges in Canada, with approximately 60% of the nation’s contaminated sites involving petroleum-related pollution (CCME, 2008). This widespread contamination, stemming from decades of industrial activity including oil and gas extraction, refining, and transportation, poses significant ecotoxicological risks to both terrestrial and aquatic ecosystems. The Canadian boreal ecozone, encompassing approximately 2.9 million square kilometers and representing one of the world’s largest intact forest ecosystems, faces particularly acute exposure to PHC contamination (Brandt, 2009). This vast region, which supports critical ecological functions including carbon sequestration, water filtration, and biodiversity conservation, experiences ongoing industrial pressures from mining, oil sand production, forestry, and oil and gas operations. The inherent vulnerability of boreal soils—characterized by their naturally acidic pH, low nutrient content, and high sensitivity to acid precipitation—compounds the ecotoxicity of PHCs on native species (Sponseller et al., 2016; Roy et al., 2023). Moreover, the region’s naturally slow rates of decomposition and ecosystem recovery make PHC contamination particularly persistent and problematic in northern environments.

Phytoremediation is a promising, cost-effective alternative to conventional excavation and disposal methods for addressing PHC contamination in boreal ecosystems (Robertson et al., 2007). This green technology leverages the natural capabilities of plants and their associated microbial communities to degrade, extract, contain, or immobilize soil contaminants. The primary mechanisms underlying PHC phytoremediation include rhizodegradation, where enhanced microbial activity in the rhizosphere facilitates contaminant breakdown (Ali et al., 2024). However, rhizodegradation effectiveness can be significantly limited by poor plant establishment and growth in contaminated soils, reduced bioavailability of target contaminants, and the inherent complexity of petroleum hydrocarbon mixtures (Robertson et al., 2007; Dagher et al., 2020).

Recent advances in understanding plant-microbe interactions have revealed the critical role of beneficial soil microorganisms in enhancing phytoremediation outcomes. Arbuscular mycorrhizal fungi (AMF) form symbiotic associations with approximately 80-90% of vascular plant species and can constitute up to 50% of total soil microbial biomass (Lanfranco et al., 2016). These fungi extend the plant root system through extensive hyphal networks, significantly expanding the rhizosphere and creating the ‘hyphosphere’—a zone of enhanced microbial activity that facilitates both nutrient acquisition and contaminant degradation.

Within this hyphosphere, AMF orchestrate sophisticated nutrient exchange networks where they receive up to 20% of plant-fixed carbon in return for mobilizing and transporting essential nutrients such as nitrogen and phosphorus to their host plants. This carbon-for-nutrient exchange creates energy-rich microsites around hyphal surfaces that stimulate the growth and metabolic activity of hydrocarbon-degrading bacteria, effectively establishing the hyphosphere as a ’carbon bridge’ between plants and symbiotic soil microorganisms (Nanjundappa et al., 2019). The enhanced nutrient cycling within the hyphosphere not only supports increased microbial biomass and enzymatic activity necessary for PHC breakdown, but also enables plants to maintain growth and physiological function under PHC stress, thereby sustaining the rhizodegradation process.

Plant growth-promoting rhizobacteria (PGPR), including species such as *Bacillus subtilis*, have also demonstrated remarkable potential for enhancing plant tolerance to PHC-contaminated soils (Ali et al., 2024). These bacteria reduce plant stress through mechanisms including the production of phytohormones, reduction of stress ethylene levels via ACC deaminase activity, and direct biodegradation of petroleum compounds (Gowtham et al., 2020). Phytohormone production represents a primary mechanism by which *B. subtilis* enhances plant growth under contaminated conditions, with strains capable of synthesizing phytohormones - indole-3-acetic acid (IAA), cytokinins, gibberellins, and abscisic acid - that stimulate plant root development, promote cell division and elongation, and improve nutrient uptake capacity. The production of ACC deaminase by *B. subtilis* and AMF play a critical role in plant stress mitigation by cleaving 1-aminocyclopropane-1-carboxylic acid (ACC), the immediate precursor to ethylene, which inhibits root elongation, promote premature senescence, and reduces overall plant vigor under PHC exposure (Gamalero and Glick, 2015).

Additionally, *B. subtilis* strains demonstrate direct PHC biodegradation capabilities through the production of key enzymes including alkane hydroxylase and alcohol dehydrogenase (Ali et al. 2024). *B. subtilis* also produces biosurfactants such as lipopeptides – surfactin (primary), fengycin, and iturin - that increase PHC bioavailability by reducing surface tension and facilitating microbial access to the hydrophobic contaminants. Biosurfactant-producing *B. subtilis* strains have been shown to boost hydrocarbon biodegradation rates by 60–90% and achieve >80% removal within one week under optimal laboratory conditions (Vázquez Rosas Landa et al., 2023). *Bacillus subtilis* ATCC 21332 is a strain of biosurfactant-producing *B. subtilis* that can reduce the surface tension of water (72 mN/m) to 26 mN/m at room temperature (Roy et al., 2025).

This study fills a critical knowledge gap regarding the AMF-PGPR enhanced rhizodegradation in boreal soils. It is also the first to investigate plant health in AMF-PGPR amended soils that are concentrated in PHCs (>10,000 mg/kg total petroleum hydrocarbons - TPH). The combined effects of AMF and the PGPR, *Bacillus subtilis* ATCC 21332, inoculation on rhizodegradation was investigated in soils containing 12,600 mg/kg and 25,700 mg/kg TPH. To maximize both remediation efficacy and ecological compatibility, native and naturalized PHC-tolerant perennials were selected. The tufted C₄ prairie grasses *Andropogon gerardii* (big bluestem) and *Panicum virgatum* (switchgrass) both produce extensive laterally spreading fibrous root systems, while *Bouteloua curtipendula* (sideoats grama) combines a fibrous tufted roots with short rhizomes and stolons to rapidly colonize soil (Wernerehl and Givnish, 2025). These species were selected as they have previously facilitated PHC rhizodegradation in the 10,000–25,000 mg/kg TPH range (McIntosh et al. 2016; Thomas et al. 2017). The eastern boreal keystone conifer *Picea mariana* (black spruce), which dominates the field site, and the forb, *Achillea millefolium* (common yarrow), whose aromatic, finely branched roots remain active in soils with 10,000–50,000 mg/kg TPH were also selected (Roy et al., 2023).

Currently, most studies examine AMF-PGPR enhanced phytoremediation over a short (<120 days) time period (Dagher et al., 2020; Lopez-Echartea et al., 2020). A 24-month timeframe was selected for this study to evaluate the long-term effects of PHC contamination, AMF, and *B. subtilis* ATCC 21332 on plant establishment. Soil microbial community dynamics during PHC rhizodegradation was investigated in a previous study, where PHC contamination was identified as the primary driver of rhizobacterial biodiversity over plant species and soil treatment (Roy et al., 2025). These insights advance sustainable, cost-effective bioremediation approaches tailored to boreal environments, offering a framework for simultaneously optimizing plant health and remediation of PHCs. The most effective remediation mechanism identified in this study may be used in conjunction with the most PHC-resilient plant species in field trials at PHC-contaminated sites across the Canadian boreal ecozone.

## 2.0 Materials and methods

### 2.1 Site soils

#### 2.11 Soil collection and preparation

As reported by Roy et al., (2025), soil samples representing background conditions (<120 mg/kg total petroleum hydrocarbons [TPH]) and PHC contamination (25, 700 mg/kg TPH) were obtained in 2022 from a former petroleum distribution terminal located in northern Ontario. This site, which stored No. 2 diesel fuel, ceased operations 25+ years ago and experienced extensive natural weathering; no refining ever took place there. Earlier investigations by Roy et al., (2023, 2024) examined soils from the contaminated area (11,900 –12,800 mg/kg TPH) and documented pronounced toxicity to indigenous plants and soil invertebrates. Both background and contaminated soils were excavated from 6 m² plots to a depth of 60 cm—with the uppermost 10 cm removed to eliminate plant material—using an excavator. A total of 42 steel drums (200 L each; 21 per soil type) were filled and then shipped to the Royal Military College of Canada (RMC), where they were sieved through a 6.25 mm soil sifter box in preparation for the greenhouse phytoremediation experiments. Sieved soils were potted into DCN Harmony Planters (30.5 cm × 30.5 cm × 45.6 cm), with each pot holding 25 L. To achieve a lower hydrocarbon level, a 1:1 blend of background and contaminated soils was prepared, resulting in a mixture containing approximately 12,600 mg/kg TPH, as described in Roy et al., (2025).

#### 2.12 Particle size and nutrients

Particle size of the soils were measured at ALS Environmental using the ASTM D6913 reference method (ASTM, 2014), as reported in Roy et al., (2025). The <120 mg/kg TPH, 12,600 mg/kg TPH, and 25,700 mg/kg TPH soils are all considered coarse-grained (CCME 2008). At the start of the greenhouse experiments, soil nutrient analyses were performed by the University of Guelph Laboratory Services, as described by Roy et al., (2023, 2024). The amount of extractable nitrogen from ammonium (ammonium-N), from nitrate (nitrate-N), from nitrite (nitrite-N), as well as phosphorous (P), potassium (K), magnesium (Mg), sodium (Na), calcium (Ca), total carbon, organic carbon, inorganic carbon, and sodium adsorption ratio (SAR) were measured in the background and contaminated site soils (Roy et al., 2025 Table 1). The <120 mg/kg TPH soil contains 4x, 7x, 1.5x, 1.5x, and 2.0x higher Ammonium-N, nitrate-N, nitrite-N, P, and Ca compared to the 25,700 mg/kg TPH soil, respectively. The 25,700 mg/kg TPH soil contains 5x higher total carbon, including 5x and 6x higher organic and inorganic carbon, compared to the <120 mg/kg TPH soil. The pH of the <120 mg/kg TPH and 25,700 mg/kg TPH soils are 7.40 and 6.40, respectively.

**Table 1.**
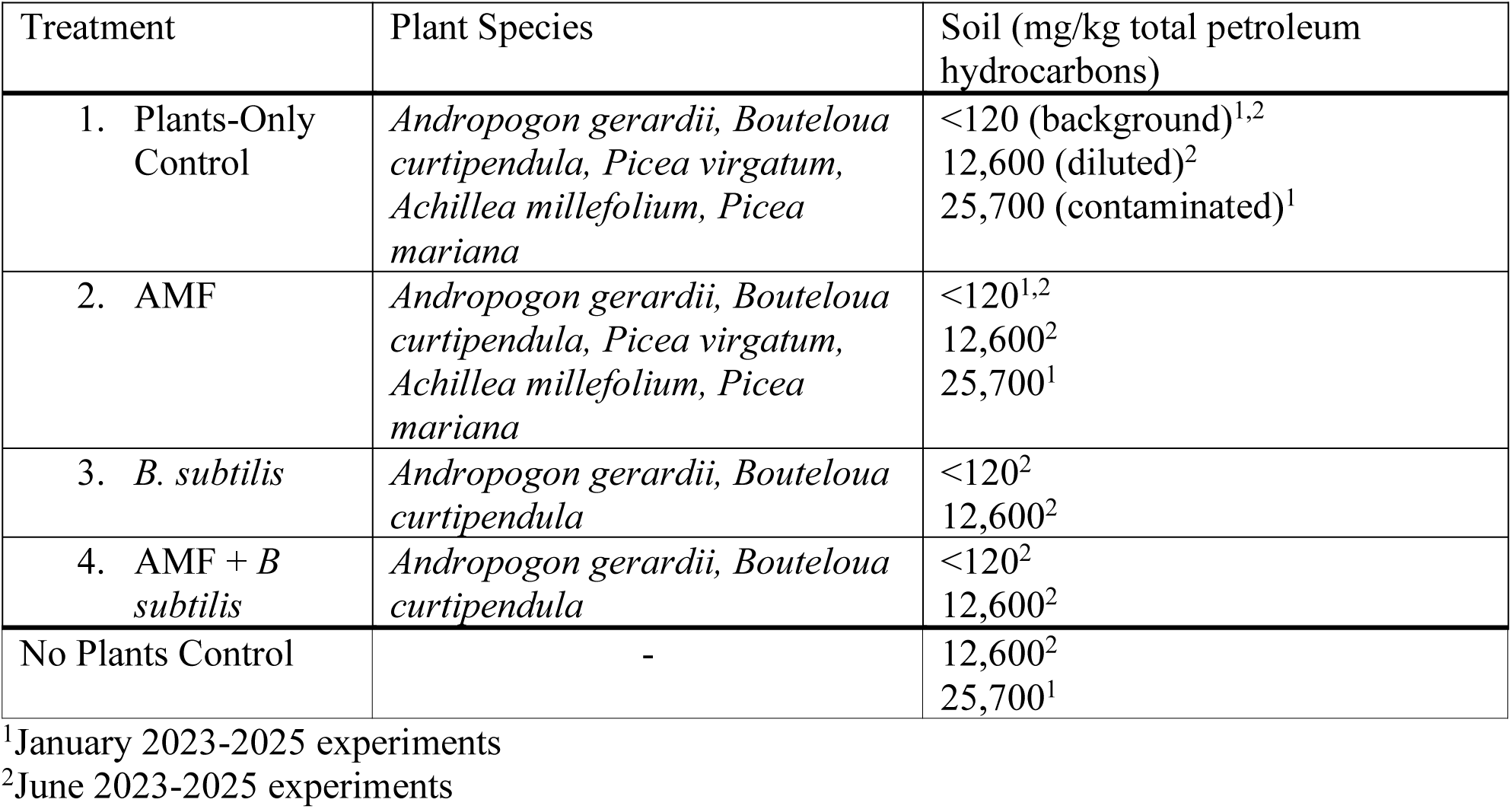
Experimental design of the 24-month greenhouse rhizodegradation experiments: January 2023-2025^1^ and June 2023-2025^2^. Four treatments were employed: 1. Plants-Only Control, 2. AMF, 3. *B. subtilis*, and 4. AMF+*B. subtilis* soil inoculants. A No Plants Control was also employed. Five replicates (*n*=5) were used for each treatment, plant species and soil.

| Treatment | Plant Species | Soil (mg/kg total petroleum hydrocarbons) |
| --- | --- | --- |
| 1. Plants-Only Control | *Andropogon gerardii*, *Bouteloua curtipendula*, *Picea virgatum*, *Achillea millefolium*, *Picea mariana* | <120 (background)<sup>1,2</sup><br>12,600 (diluted)<sup>2</sup><br>25,700 (contaminated)<sup>1</sup> |
| 2. AMF | *Andropogon gerardii*, *Bouteloua curtipendula*, *Picea virgatum*, *Achillea millefolium*, *Picea mariana* | <120<sup>1,2</sup><br>12,600<sup>2</sup><br>25,700<sup>1</sup> |
| 3. *B. subtilis* | *Andropogon gerardii*, *Bouteloua curtipendula* | <120<sup>2</sup><br>12,600<sup>2</sup> |
| 4. AMF + *B. subtilis* | *Andropogon gerardii*, *Bouteloua curtipendula* | <120<sup>2</sup><br>12,600<sup>2</sup> |
| No Plants Control | - | 12,600<sup>2</sup><br>25,700<sup>1</sup> |

<sup>1</sup>January 2023-2025 experiments
<sup>2</sup>June 2023-2025 experiments

#### 2.13 Initial PHC concentrations

Canadian Council Ministers of the Environment (CCME) Fractions (F)2-4 concentrations (F2: >C10-C16, F3: >C16-C32; F4: >C32-C50 effective carbon numbers) were quantified in December 2022. Fraction 1 concentrations were not measured in this study due to below-detection limit levels (<10 mg/kg) during preliminary quantification (Roy et al., 2025). Analysis was conducted at the CALA-accredited Analytical Services Unit, Queen’s University, employing the CCME reference method (CCME, 2008). Three barrels were randomly selected for each soil type, with three samples analyzed per barrel. The analytical procedure involved 1:1 hexane: acetone solvent extraction, ultrasonication, rotary evaporation using a Büchi rotovapor R-114, silica gel cleanup, and gas chromatography with flame ionization detection (Agilent GC 7890A). Subsequently, in July 2023, CCME F2-F4 concentrations were determined in the 12,600 mg/kg TPH soil samples collected from six unplanted greenhouse pots using identical analytical protocols. Quality control measures included one method blank, one sample duplicate, and a 5,000 mg/kg diesel fuel quality control spike for each PHC analysis.

Both site soils underwent metal contamination screening at the Analytical Services Unit following US EPA Method 200.7 (US EPA, 1994). Sample preparation involved grinding and overnight drying of 0.5 g soil samples, followed by aqua regia digestion on a DigiPREP LS graphite block (SCP Science) for four hours at 95°C. Digested samples were analyzed using inductively-coupled plasma-optical emission spectroscopy (Thermo iCAP 7400 Dual view spectrometer). The analysis confirmed the absence of metal contamination in both soil types (Roy et al., 2025)

### 2.2 Greenhouse experiments

#### 2.21 Plant seeds

Seeds of *A. gerardii*, *B. curtipendula*, *A. millefolium*, and *P. virgatum* were procured from Sheffield’s Seed Company in 2022, with corresponding lot numbers 1834306, 1832814, 1833869, and 1834252, respectively. *P. mariana* seeds were shipped to RMC from the Saskatchewan Research Council in 2015. Germination viability testing was conducted for each species using parafilm-sealed Petri dishes containing 10 g of background site soil as the growth medium, as described in Roy et al., (2025). All tested species demonstrated high viability, with germination rates exceeding 80% across all Petri dishes, confirming the suitability of the selected seed lots for experimental use.

#### 2.22 Experimental setup

In January 2023, 24-month greenhouse phytoremediation experiments were initiated at RMC to evaluate the effectiveness of selected soil treatments on remediation of PHC-contaminated field soil (Table 1). The five selected plants were used in conjunction with treatment treatments in the <120 mg/kg (background) and 25,700 mg/kg TPH (contaminated) soils: 1. Plants-Only Control and 2. arbuscular mycorrhizal fungi (AMF) soil treatment. In June 2024, the two best growing plant species - *A. gerardii* and *B. curtipendula* – were selected for phytoremediation trials in the background and 12,600 mg/kg TPH soils, under four treatments: 1. Plants-Only Control, 2. AMF, 3. *Bacillus subtilis* (ATCC 21332), and 4. AMF+*B. subtilis*.

Each AMF and AMF+*B. subtilis* pot was inoculated with 13.2 g of with MycoApply Ultra Fine Endo © for the AMF treatment, comprising of the four Glomeromycota fungi: *Rhizophagus irregularis* (formerly *Glomus intraradices*)*, Glomus mosseae, G. aggregatum,* and *G. etunicatum*. A total of 42 mL of active *B. subtilis* ATCC 21332 culture was added to each *B. subtilis* and AMF+*B. subtilis* treatment pot (mean absorbance at 600 nm: 1.87; cell dry weight: 4 g/L). The methodology for culturing and enumerating *B. subtilis* ATCC 21332 is described in Roy et al., (2025). A No Plants Control was also employed for both the January 2023-2025 and June 2023-2025 experiments (Table 1).

#### 2.23 Seed counts

Prior to sowing, *P. mariana* seeds underwent a six-week cold stratification in *Sphagnum* peat moss and deionized water to enhance germination. The seeds of the remaining four species were subjected to a 24-hour hydration period for scarification. All seeds were evenly distributed within the upper 1-cm of soil. During the January 2023-2025 experiments, seeding rates for all species except *P. mariana*, were set at 20% above the pure live seed count, as detailed in Roy et al., (2025). Seeding counts per pot were as follows: 41 – *A. gerardii*, 16 – *B. curtipendula*, 53 – *P. virgatum*, 61 – *A. millefolium*, and 12 – *P. mariana*.

#### 2.24 Greenhouse settings

The greenhouse facility at RMC was maintained at temperatures ranging from 20°C in winter to 30°C in summer. Temperature regulation was facilitated by an iGrow 1800 Advanced Environmental Controller operating with the LinkConn 1000 v3.6 software. During the winter months, light intensity was maintained between 260 and 660 W/m², and between 780 and 1120 W/m² during summer. Natural sunlight was used as the primary light source; however, when light intensity fell below 350 W/m², supplemental illumination was provided using Verilux Full Spectrum H588 light rods. Relative humidity was consistently maintained at 50% year-round. Pots were irrigated three times per week, with each pot receiving approximately 3 liters of water, weekly. Manual removal of weeds and non-target plant species was performed as needed.

### 2.3 Plant harvest

#### 2.31 Above- and below-ground biomasses

Above- and below-ground plant-fungi biomass measurements were collected following the 24-month experimental periods to establish plant health in response to PHCs (12,600 mg/kg and 25,700 mg/kg TPH) and soil treatments. During the January 2023-2025 experiments, plants were watered consistently until harvest in January 2025. This facilitated collection of wet and dry above- and below-ground plant biomass measurements and percent moisture measurements.

Plants were not watered for three weeks prior to the June 2023-2025 harvest to facilitate the below-ground biomass handling process. The plants were fully dried during the June 2025 harvest.

USDA National Resource Conservation Service methods were referenced to harvest biomasses (USDA, 2022): drain-spades (41 cm length x 14 cm width) purchased from Grainger Canada were used to loosen the soil around the plants, non-stick micro-tip pruning and titanium snips were used for separating above- and below-ground biomasses, and 710 μm - 8 mm diameter mesh sieves (McMaster Carr) were used to filter the below-ground biomasses before and after drying. Pre-weighed aluminum steam pans (U-Line) were used to hold the biomasses. Wet biomasses were dried in a Kratos Full-Size Non-Insulated Holding and Proofing Cabinet (Hubert Canada) at 70 °C for 48 hours (January 2023-2025 experiments). Percent moisture in the above- and below-ground biomasses were calculated as follows:

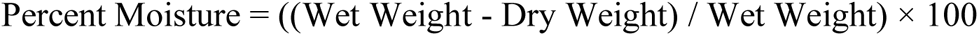

Above- and below-ground biomasses for all species, except *P. mariana*, were weighed on a Ohaus 30253019 Scout Pro balance (0.1 g – 6,000 g range). *P. mariana* was weighed on a Denver Instrument XP-300 balance for enhanced precision (0.01- 300 g range). The weight of the aluminum pans were subtracted from each biomass measurement.

#### 2.32 Plant height

Above-ground length measurements were taken from five randomly selected shoots in each pot (*n* = 25), every three to six months throughout the experiments as a non-invasive preliminary indicator of plant growth. All plants grew consistently over the course of the experiments. Following the January and June 2025 plant harvests, the longest above-ground and below-ground length measurements were taken from each treatment pot (*n*= 5). All plants, except *P. mariana* in the January 2023-2025 experiments, reached the full length of the pots (46 cm) following two years of growth – across all treatments. Therefore, below-ground length measurements were not analyzed further. *P. mariana* reached a mean (± SD) below-ground length measurement of 0.60 (0.20) and 1.20 (0.50) cm in the Plants-Only Control and AMF treatments in the <120 mg/kg TPH background soil, respectively. Mean below-ground lengths of 13.2 (2.0) and 18.8 (2.3) cm were respectively reached in the 25,700 mg/kg TPH treatment soil treatments.

### 2.4 Final PHC measurements

Final PHC concentrations for the January 2023 – 2025 experiments were measured in each treatment pot in January of 2025 (Fig. 1). Soil PHC measurements for each treatment pot (*n*= 5) was taken for the June 2023 – 2025 experiments in December 2024, 17 months into the experiments, with the intention of remeasuring final PHC measurements in June 2025. However, PHC concentrations were remediated to federal CCME and provincial guidelines within the 17 month period (Fig. 2). Hence, 24-month soil PHC concentration measurements were not taken.

**Fig. 1.**
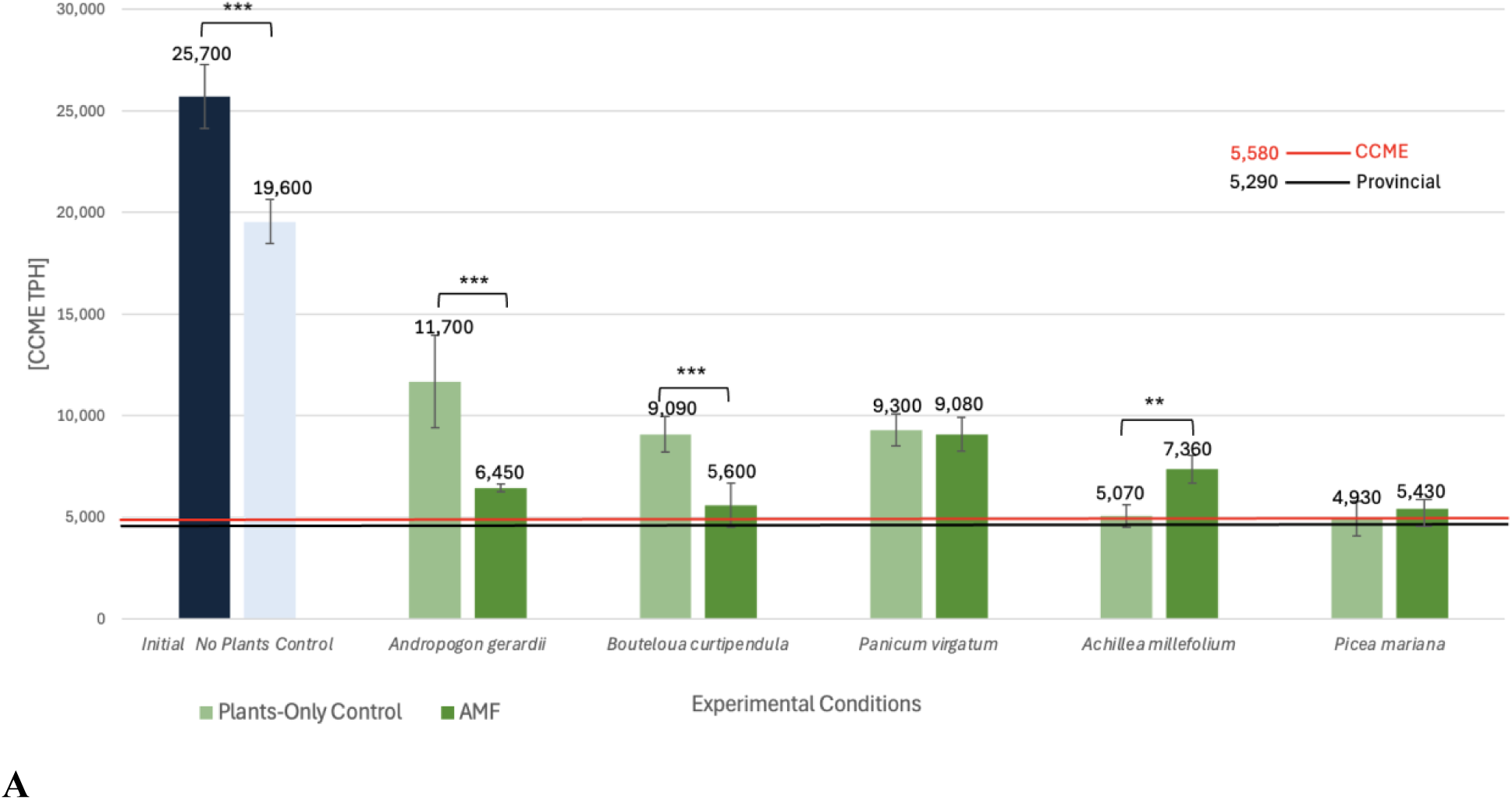
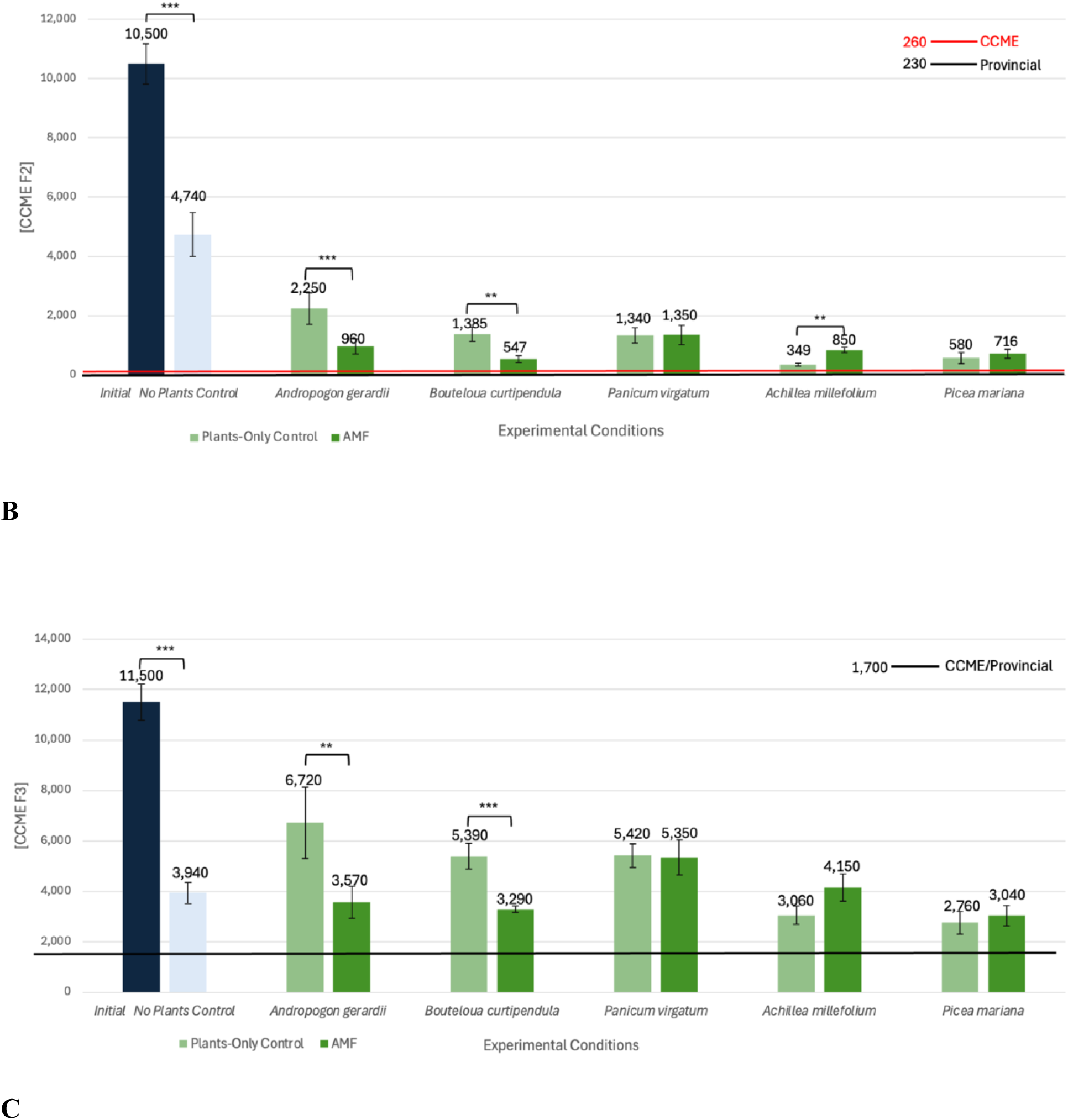
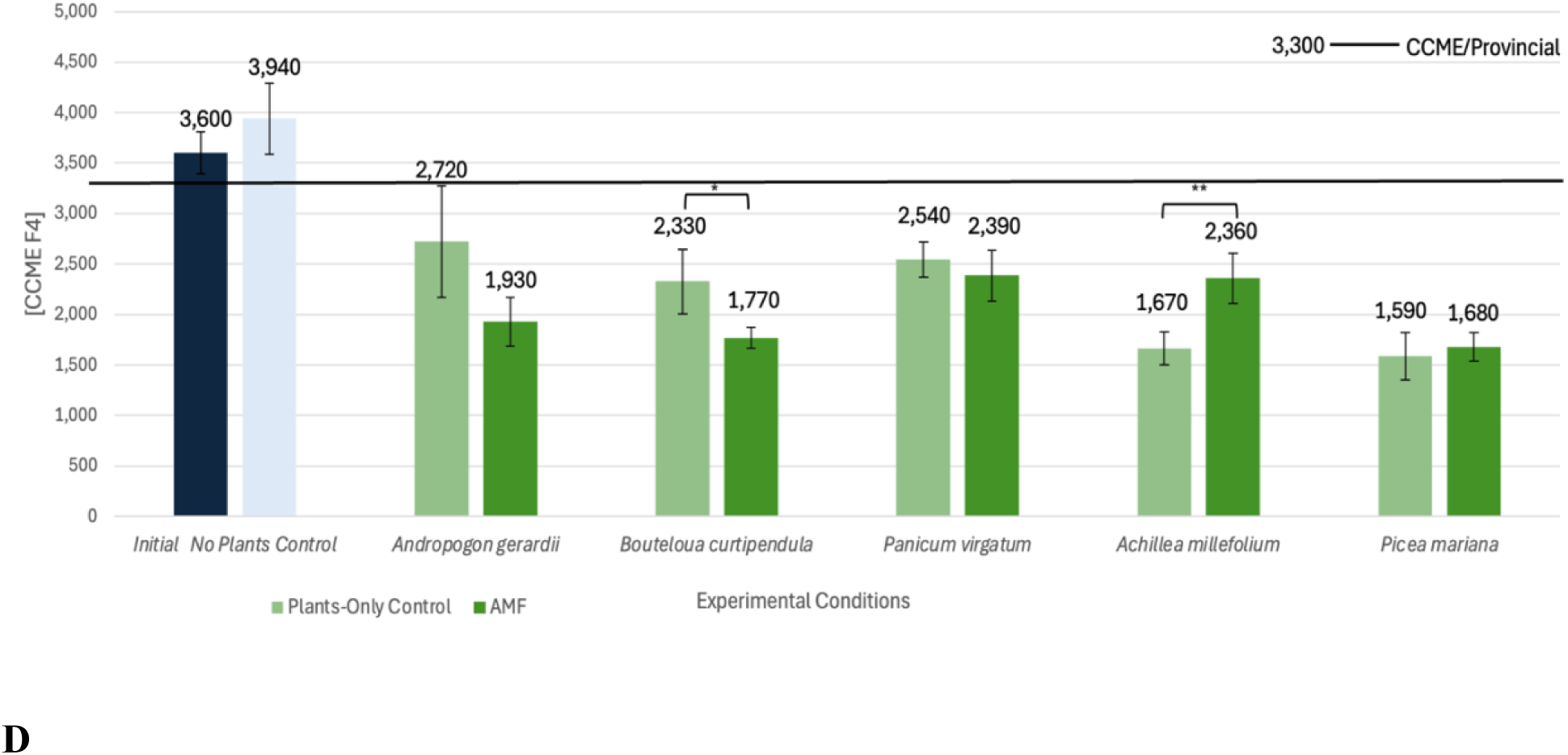
Barplots of Canadian Council of Ministers of the Environment (CCME) soil PHC concentrations in the soil during the January 2023-2025 experiments. The initial PHC concentration is indicated by the dark blue bar. Remediation in the No Plants Control is indicated by the light blue bars. Phytoremediation under the Plants-Only Control (light green) and AMF (dark green) treatment regimens are indicated with the green bars. Standard error is indicated by the error bars. Horizontal red and black lines indicate federal CCME and Ontario provincial guidelines (CCME 2008; OME 2011). Mean concentrations are written above the error bars. Brackets above bars indicate significant pairwise Mann-Whitney *U* test between the initial concentration and No Plants Control and between the phytoremediation treatments (*p* < 0.001 = "***", *p* < 0.01 = "**", *p* < 0.05 = "*"). **A.** CCME TPH concentrations. **B.** CCME F2 concentrations. **C.** CCME F3 concentrations. **D.** CCME F4 concentrations.

**Fig. 2.**
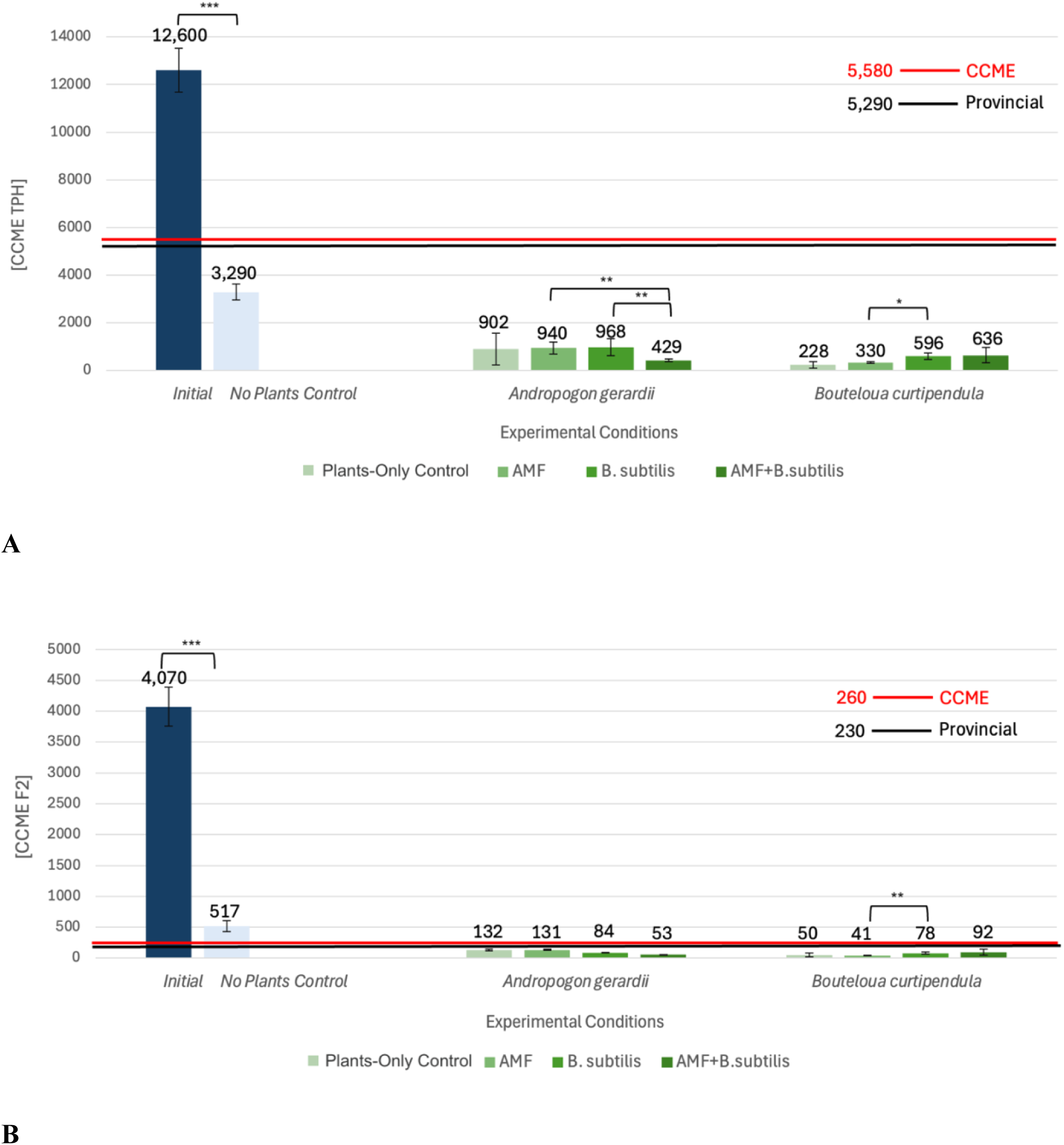
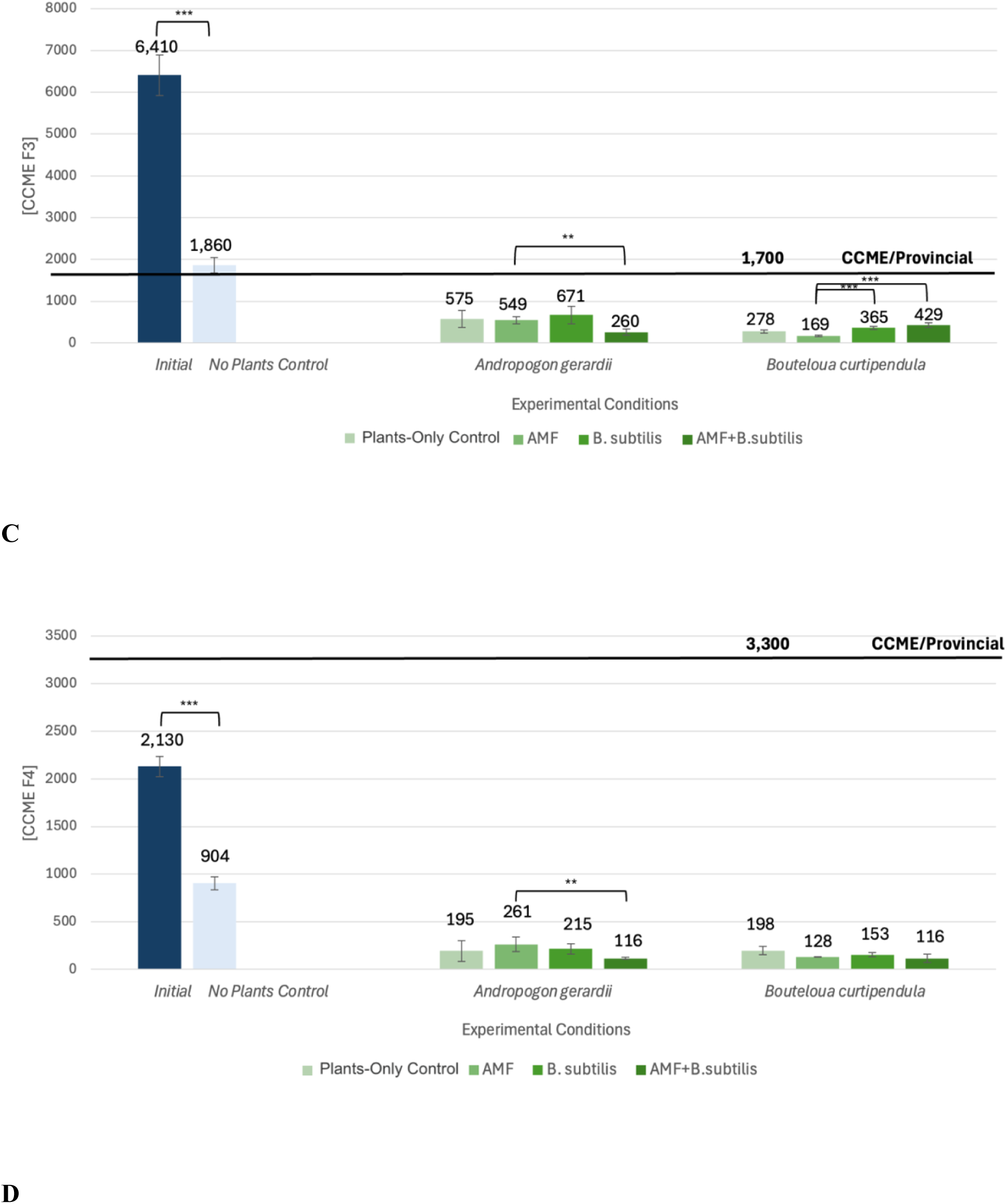
Barplots of Canadian Council of Ministers of the Environment (CCME) soil PHC concentrations during the June 2023-2025 experiments. The initial PHC concentration is indicated by the dark blue bar. Remediation in the No Plants Control is indicated by the light blue bars. Phytoremediation under the Plants-Only Control, AMF, *B. subtilis*, and AMF+*B. subtilis* treatment regimens are indicated on a light (Plants-Only Control) to dark green (AMF+*B. subtilis*) colour gradient. Standard error is indicated by the error bars. Mean concentrations are written above the error bars. Horizontal red and black lines indicate federal CCME and Ontario provincial guidelines (CCME 2008; OME 2011). Brackets above bars indicate significant pairwise Mann-Whitney *U* test comparisons between the initial concentration and No Plants Control and between the phytoremediation treatments (*p* < 0.001 = "***", *p* < 0.01 = "**", *p* < 0.05 = "*"). **A.** CCME TPH concentrations. **B.** CCME F2 concentrations. **C.** CCME F3 concentrations. **D.** CCME F4 concentrations.

### 2.5 Theory and Calculations

All statistical data was analyzed in RStudio 2024.12.1563 using the “vegan”, “dplyr”, “ggplot2”, “effsize”, “ggsignif”, “tidyr”, and “rstatix” libraries. Figures (Figs.) 1-2 were created in Microsoft Excel for Mac Version 16.96. Subsequently figures (Figs. 3-10) were created in RStudio 2024.12.1563. Two-tailed Mann-Whitney *U* tests with Bonferroni corrections were used for pairwise statistical comparisons (*n* ≥ 5) and Hedge’s *g* for corresponding effect size comparisons. Hedge’s *g* corrects for small sample bias (*n* < 20) and is based on the Cohen’s *d* formula. Absolute values (*g*) of |0.15|, |0.40|, and |0.75| were considered thresholds of small, medium, and large effect size (Brydges, 2019). Two-tailed Kruskal-Wallis Tests and eta-squared effect sizes (η²H) were employed when three or more independent groups were compared. Eta-squared (η²H) of 0.01, 0.06, and ≥ 0.14 indicating small, moderate, and large effects, respectively (Richardson, 2011). Outliers were not removed from the dataset to facilitate sufficient replication for significance testing.

**Fig. 3.**
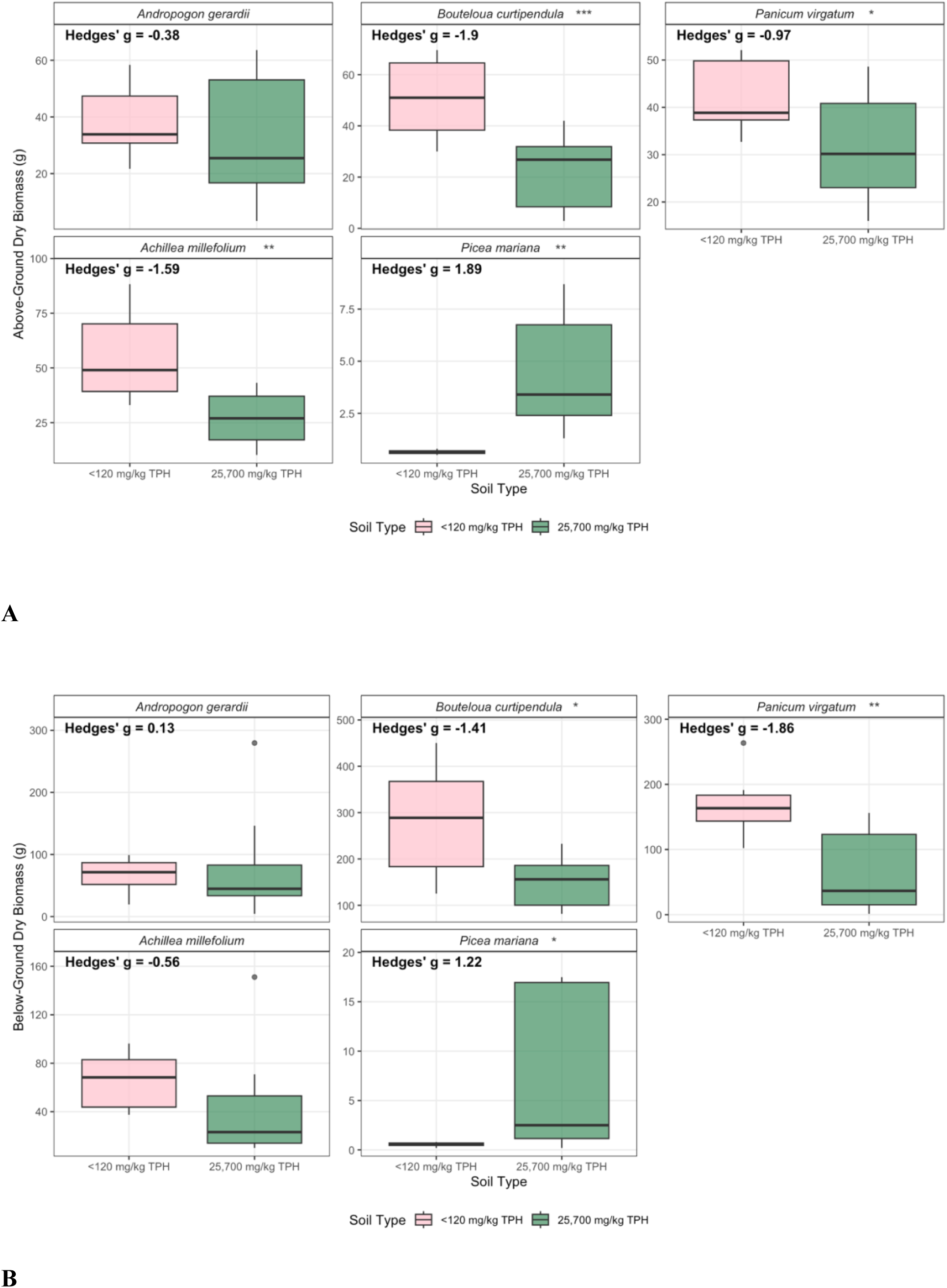
Boxplots of dry biomass (g) of the five plant species that grew in the <120 mg/kg TPH and 25,700 mg/kg TPH site soils. Mann-Whitney *U* test significant pair-wise comparisons are represented by asterisks beside plant species names in each facet (*p* < 0.001 = "***", *p* < 0.01 = "**", *p* < 0.05 = "*"). Hedge’s *g* indicate effect size comparisons between the site soils. **A**. Above-ground biomass. **B**. Below-ground biomass. AMF inoculation significantly enhanced above-ground dry biomass in both the <120 mg/kg TPH and 25,700 mg/kg TPH soils (Mann Whitney *U* tests: both *p* < 0.001), with very large effects (Hedge’s *g* = 1.09 and 1.60) relative to the Plants-Only Control (Fig. 4). Below-ground responses diverged by PHC soil contamination; AMF increased below-ground biomass moderately in the <120 mg/kg TPH soil (*p* = 0.05; *g* = 0.58) but produced a large, significant enhancement in the 25,700 mg/kg TPH soil (*p* < 0.001; *g* = 1.37). AMF significantly improved both above-ground biomass and below-ground biomass in all plant species in the 25,700 mg/kg TPH soil (all *p* < 0.05), with large effects in both above-ground (*g* = 1.55 to 2.06) and below-ground dry biomass (*g* = 1.37 to 2.85).

**Fig. 4.**
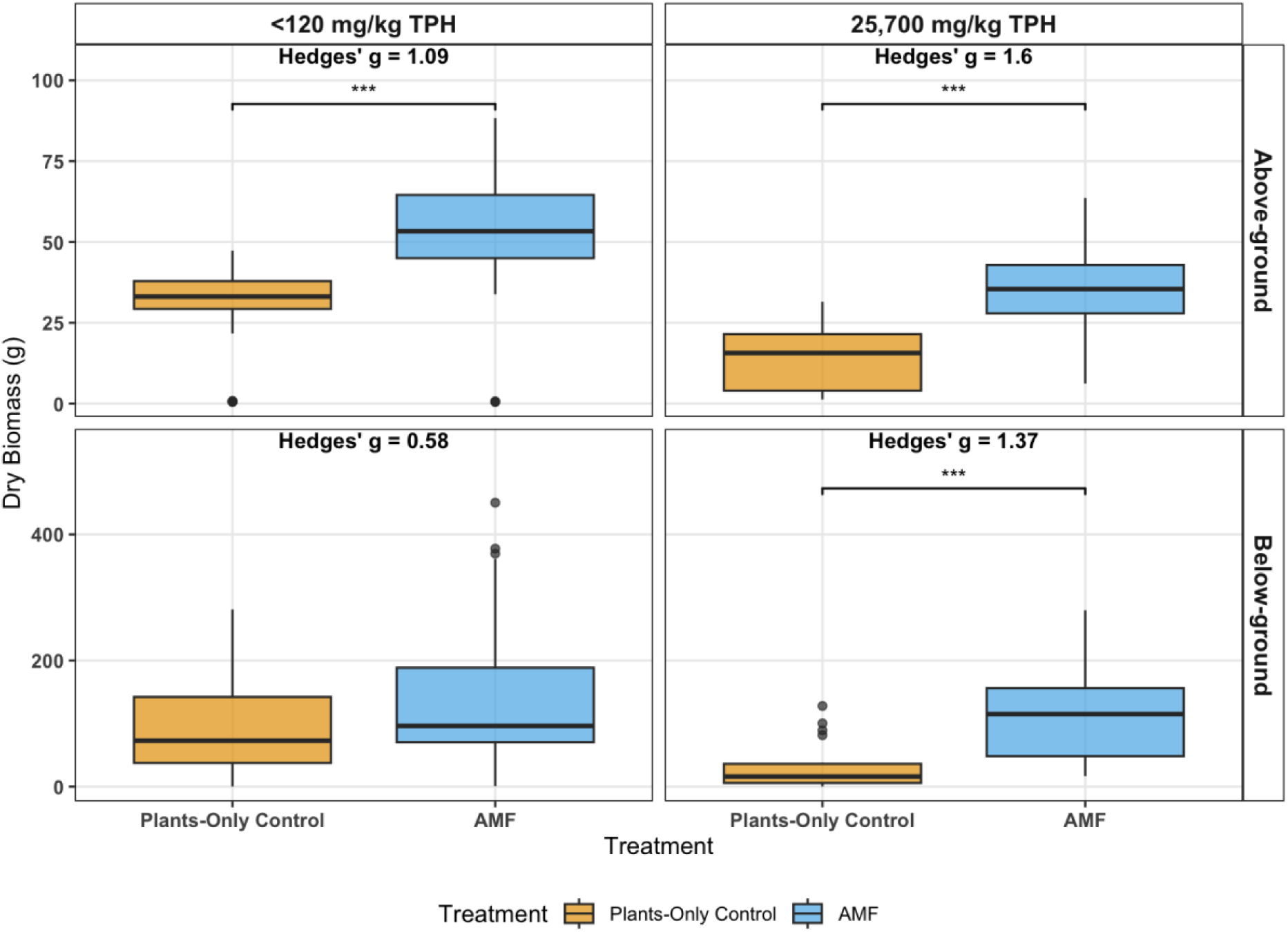
Boxplots of above- and below-ground dry biomass (g) in the Plants-Only Control and AMF treatments. Boxplots are faceted by above-ground (top facets) and below-ground (bottom facets) compartments in the <120 mg/kg TPH (left facets) and 25,700 mg/kg TPH (right facets) soils. Brackets and asterisks above boxplots indicate significant Mann-Whitney *U* test pairwise comparisons between treatments (*p* < 0.001 = "***", *p* < 0.01 = "**", *p* < 0.05 = "*"). Hedge’s *g* indicate effect size comparisons between treatments.

The ratio of below-ground to above-ground dry biomasses and below-ground to above-ground moisture content are referred to as “root: shoot” ratios. The root: shoot ratios facilitate interpretation of plant resource allocation as studies have suggested that plants may allocate more resources to root growth in response to environmental stressors such as PHCs within the first few weeks to months of growth (Desalme et al., 2011). Multi-year plant root: shoot allocation in response to PHC stress has not been investigated in previous studies.

## 3.0 Results

### 3.1 Soil PHC remediation

Following two years of remediation, mean PHC concentrations in the 25,700 mg/kg TPH soil were significantly reduced across all phytoremediation treatment conditions (Fig. 1A; Kruskal-Wallis test: *p* = 0.003). In the No Plants Control, natural attenuation processes reduced TPH to 19,600 mg/kg, indicating a significant decrease (24% TPH removal; Mann-Whitney *U* test, *p* < 0.001) with a large effect (Hedge’s *g* = 1.98). The Plants-Only Control treatment further enhanced remediation efficacy, achieving a mean concentrations of 8,500 mg/kg TPH, indicating a statistically significant improvement over the No Plants Control (*p* = 0.004), with a larger effect (67% TPH removal; *g* = 2.19), across all five plant species. The AMF treatment yielded reduced TPH to a mean concentration of 6,780 mg/kg TPH, which was significantly lower than both the No Plants Control (*p* = 0.002) and the Plants-Only Control (*p* < 0.001), and corresponded to respective mean TPH removal rates of 74% and 65%.

Similar patterns of enhanced remediation were observed across individual CCME PHC fractions, with AMF-enhanced phytoremediation achieving significant reductions in F2 (mean removal: 92%), F3 (66%), and F4 (56%) compared to the No Plants Control (Fig. 1B–D). The No Plants Control was not able to reduce F4 levels below federal CCME or provincial guidelines (CCME 2008; OME 2011). Both the Plants-Only Control and AMF treatments brought F4 concentrations below both guideline thresholds within the 24-month period (Fig. 1D), representing an overall large effect for the Plants-Only Control (*g* = 1.51) and a larger effect for the AMF treatment (*g* = 1.59).

There was no significant difference in TPH removal across the five plant species (Kruskal–Wallis test: *p* = 0.41). However, *P. mariana* achieved the lowest final mean TPH concentration (4,930 mg/kg TPH), representing increased TPH removal of 2.8%, 46%, 47%, and 58% compared to *A. millefolium*, *B. curtipendula*, *P. virgatum*, and *A. gerardii*, respectively (Fig. 1). Pairwise Mann-Whitney *U* tests confirmed that *P. mariana* reduced TPH significantly more than *B. curtipendula*, *P. virgatum*, and *A. gerardii* (all *p* < 0.05) in the Plants-Only Control.

After 17 months of remediation, the TPH concentrations in the 12,600 mg/kg soil were reduced to below both federal CCME and Ontario provincial guideline thresholds in all planted treatments (Fig. 2A; CCME, 2008; OME, 2011). For *A. gerardii* pots, the AMF+ *B. subtilis* treatment resulted in the highest degree of TPH remediation (97% removal) while the Plants-Only Control resulted in 98% TPH remediation for *B. curtipendula* (Figs. 2A-D). In the No Plants Control, mean TPH declined significantly from 12,600 to 3,290 mg/kg (Fig. 2A; Mann– Whitney *U* test: *p* < 0.001), with a large effect (Hedge’s *g* = 2.45). The Plants-Only Control treatment further accelerated this decline, lowering TPH to a mean concentrations of 565 mg/kg across both species (*p* = 0.01), while AMF inoculation resulted in remediation to 636 mg/kg TPH, representing a non-significant difference compared to the Plants-Only Control (*p* = 0.11). *B. subtilis* and the combined AMF+*B. subtilis* treatments yielded mean TPH of 782 mg/kg and 533 mg/kg, respectively, representing non-significant decreases compared to the Plants-Only Control.

CCME F2-F4 concentrations declined to 517 (F2), 1,860 (F3), and 904 mg/kg (F4), respectively, in the No Plants Control —meeting federal and provincial guideline values for F4 but remaining above the federal/provincial F2 and F3 thresholds (Fig. 2B–D; CCME 2008; OME 2011). The Plants-Only Control reduced F3 to 575 mg/kg, meeting federal and provincial guidelines (Fig. 2C). Across both grass species, the AMF+*B. subtilis* treatment achieved the highest magnitude remediation compared to the No Plants Control, with mean removal rates of 84% (TPH), 86% (F2), 81% (F3) and 87% (F4). Compared to the initial starting concentrations, AMF+*B. subtilis* reduced mean PHC levels by 96% (TPH), 98% (F2), 95% (F3) and 95% (F4).

### 3.2 Plant biomass (January 2023 – 2025)

There were significant decreases in both above-c and below-ground dry biomass (g) in the 25,700 mg/kg TPH soil compared to the <120 mg/kg TPH soil (Fig. S1; Mann-Whitney *U* tests: both *p* < 0.05), with moderate effect size in the below-ground (*g* = - 0.51) and large effect in the above-ground biomass (*g* = - 0.72). Root: shoot biomass ratios were not significantly different between the two soils, with root growth favoured over shoot growth by two-fold in both soils (*p* = 0.27; *g* = 0.24). Species-specific responses to the 25,700 mg/kg TPH soil were observed (Fig. 3A). *A. gerardii* did not show a statistically significant decrease in above-ground biomass between the <120 and 25,700 mg/kg TPH soils (*p* = 0.38), but a moderate reduction in biomass was observed in the 25,700 mg/kg TPH soil (*g* = - 0.38). In contrast, *B. curtipendula, P. virgatum*, and *A. millefolium* exhibited significant reductions (all *p* < 0.05) in above-ground biomass in the 25,700 mg/kg TPH soil with large negative effects (g = - 0.97 to -1.90). *P. mariana* showed a significant (*p* = 0.001) and large effect (*g* = 1.89) increase in above-ground biomass in the 25,700 mg/kg TPH soil.

Although the three tufted grass species exhibited similar above-ground biomasses in the <120 mg/kg TPH soil (Fig. 3A), the median below-ground biomass of *B. curtipendula* was approximately 1.7-fold higher than that of *P. virgatum* and approximately 2.5-fold higher than that of *A. gerardii* in the < 120 mg/kg TPH soil (Fig. 3B). Median *B. curtipendula* below-ground biomass was 1.7-fold higher than both *A. gerardii* and *P. virgatum* in the 25,700 mg/kg TPH soil. *A. gerardii* and *A. millefolium* showed no significant differences between the <120 and 25,700 mg/kg TPH soils, although *A. millefolium* showed a moderate decrease in below-ground biomass in the 25,700 mg/kg TPH soil (*g* = - 0.56). *B. curtipendula* and *P. virgatum* both exhibited significant (*p* < 0.05) and large effect reductions in below-ground biomass in the 25,700 mg/kg TPH soil (*g* = - 1.41 to - 1.86). *P. mariana* showed a significant and large positive increase in below-ground biomass in the 25,700 mg/kg TPH soil (*p* = 0.001; *g* = 1.22).

### 3.3 Plant moisture (January 2023 – 2025)

Plant above-ground moisture content (%) was significantly impacted by PHC contamination, although below-ground moisture was not (Fig. S2; Mann Whitney *U* tests: *p* < 0.05). Above-ground moisture was significantly higher in the <120 mg/kg TPH soil compared to the 25,700 mg/kg TPH soil (*p* <0.001; *g* = - 0.99). In contrast, below-ground moisture did not differ significantly between the soils (*p* = 0.11; *g* = - 0.35). Comparisons of above-ground moisture content based on soil type revealed that the 25,700 mg/kg TPH soil significantly reduced above-ground moisture in the four herbaceous plant species: *A. gerardii*, *B. curtipendula*, *P. virgatum*, and *A. millefolium* (Fig. 5A). These species all exhibited significantly lower above-ground moisture in the 25,700 mg/kg TPH soil compared to the <120 mg/kg TPH soil (Mann Whitney *U* tests: all *p* < 0.05), with large negative effects (*g* = -1.55 to -3.08). In contrast, *P. mariana* showed an insignificant increase in above-ground moisture content in the 25,700 mg/kg TPH soil (*p* = 0.06; *g* = 1.40).

**Fig. 5.**
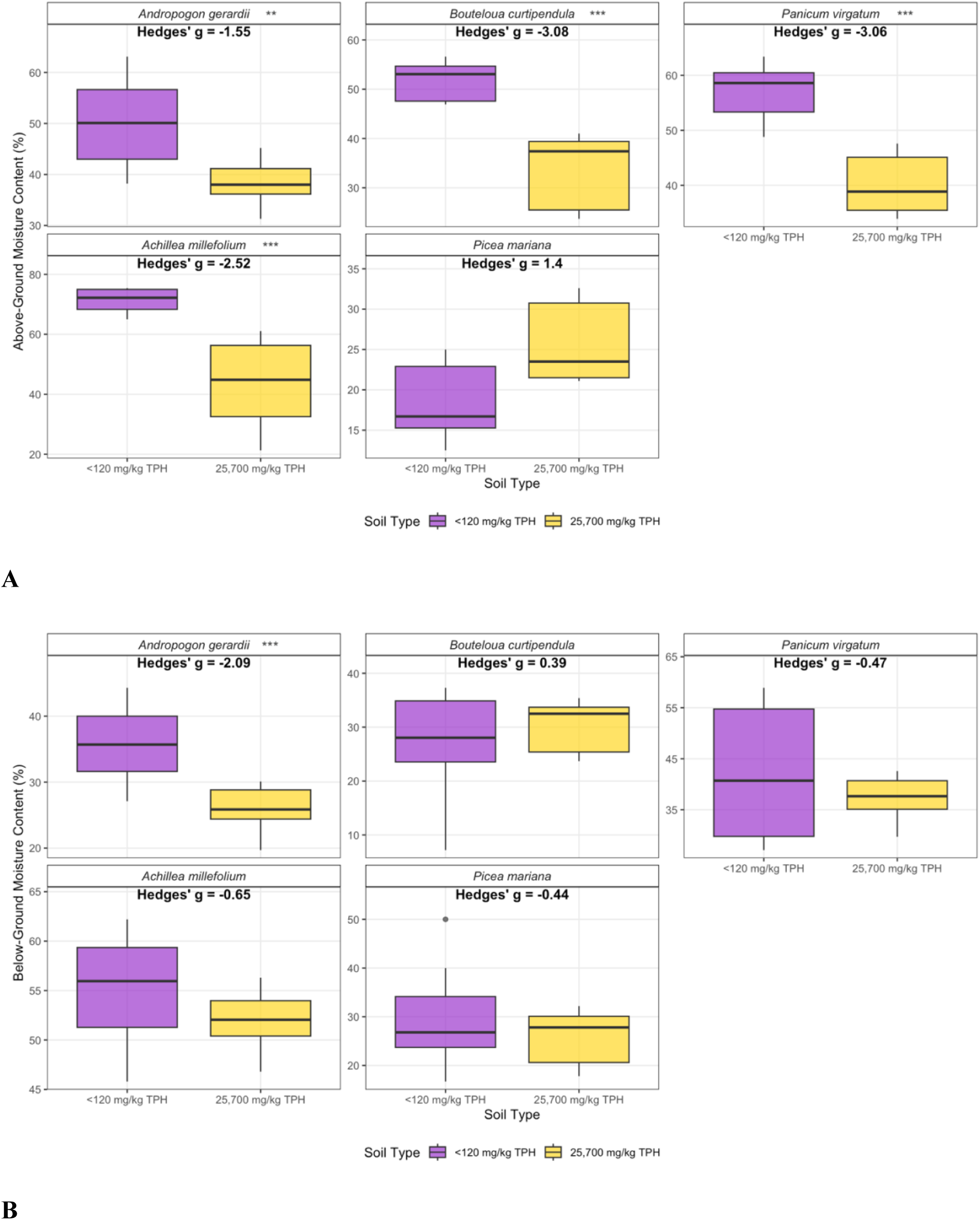
Boxplots of moisture content (%) in the five plant species in the <120 mg/kg TPH and 25,700 mg/kg TPH site soils. Mann-Whitney *U* test significant pair-wise comparisons are represented by asterisks beside plant species names in each facet (*p* < 0.001 = "***", *p* < 0.01 = "**", *p* < 0.05 = "*"). Hedge’s *g* indicate effect size comparisons between the site soils. **A**. Above-ground moisture. **B**. Below-ground moisture.

Below-ground moisture content revealed species-specific responses to the 25,700 mg/kg TPH soil (Fig. 5B). *A. gerardii* exhibited significantly lower below-ground moisture in the 25,700 mg/kg TPH soil compared to the <120 mg/kg TPH soil (*p* <0.001; *g* = -2.09). *B. curtipendula*, *P. virgatum*, *A. millefolium,* and *P. mariana* all showed no significant differences in below-ground moisture between soil types. *P. virgatum*, *A. millefolium,* and *P. mariana* showed moderate reductions in below-ground moisture in the 25,700 mg/kg TPH soil (*g* = -0.44 to -0.65), whereas *B. curtipendula* showed a moderate increase (*g* = 0.39).

Both above- and below-ground plant moisture content (%) varied significantly between the Plants-Only Control and AMF treatments in the 25,700 mg/kg TPH soil (Fig. S3; Mann-Whitney *U* tests: both *p* < 0.05). AMF enhanced above-ground plant moisture with a large effect (*g* = 0.63) and below-ground moisture with a moderate effect (*g* = 0.46). In the Plants-Only Control, the 25,700 mg/kg TPH pots exhibited a significantly higher root: shoot moisture ratio than the <120 mg/kg TPH soil (Mann-Whitney *U* test: *p* = 0.004), with moderate effect (*g* = 0.38). There was no significant difference in root: shoot moisture allocation between the <120 and 25,700 mg/kg TPH soils with the AMF inoculant (*p* = 0.16).

Above-ground plant moisture content (%) differed significantly in the 25,700 mg/kg TPH soil between the Plants-Only Control and AMF treatments in *A. gerardii* (Fig. 6A; Mann-Whitney *U* test: *p* = 0.009) with a large effect (*g* = 1.52). Below-ground moisture content varied among plant species based on treatment (Fig. 6B). The three tufted grasses showed significantly enhanced below-ground biomass under the AMF treatment with large effects (*p* < 0.05; *g* = 0.87 to 1.74). Below-ground moisture decreased insignificantly in *A. millefolium* and *P. mariana* with the AMF amendment (*g* = -0.04 to -0.37).

**Fig. 6.**
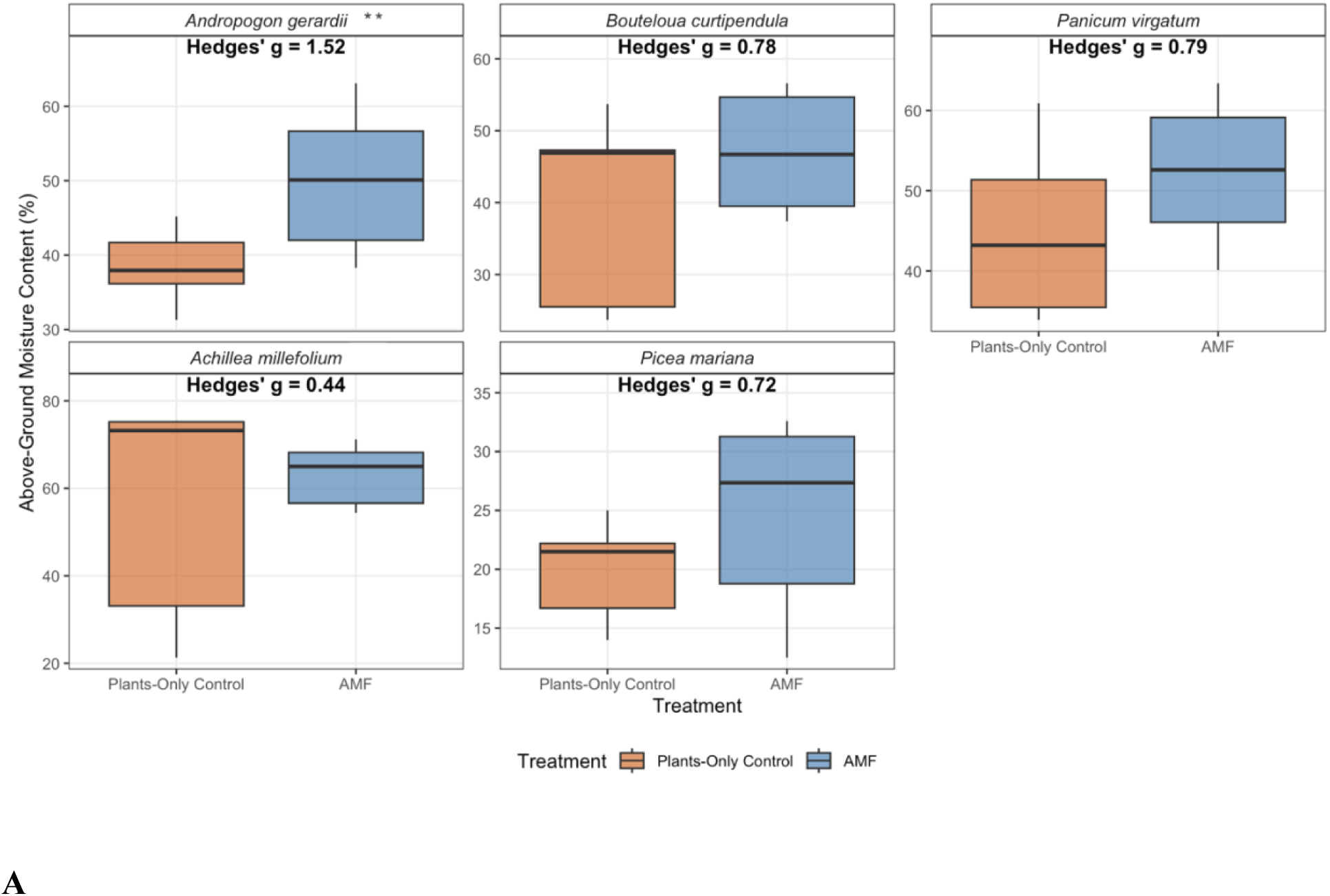
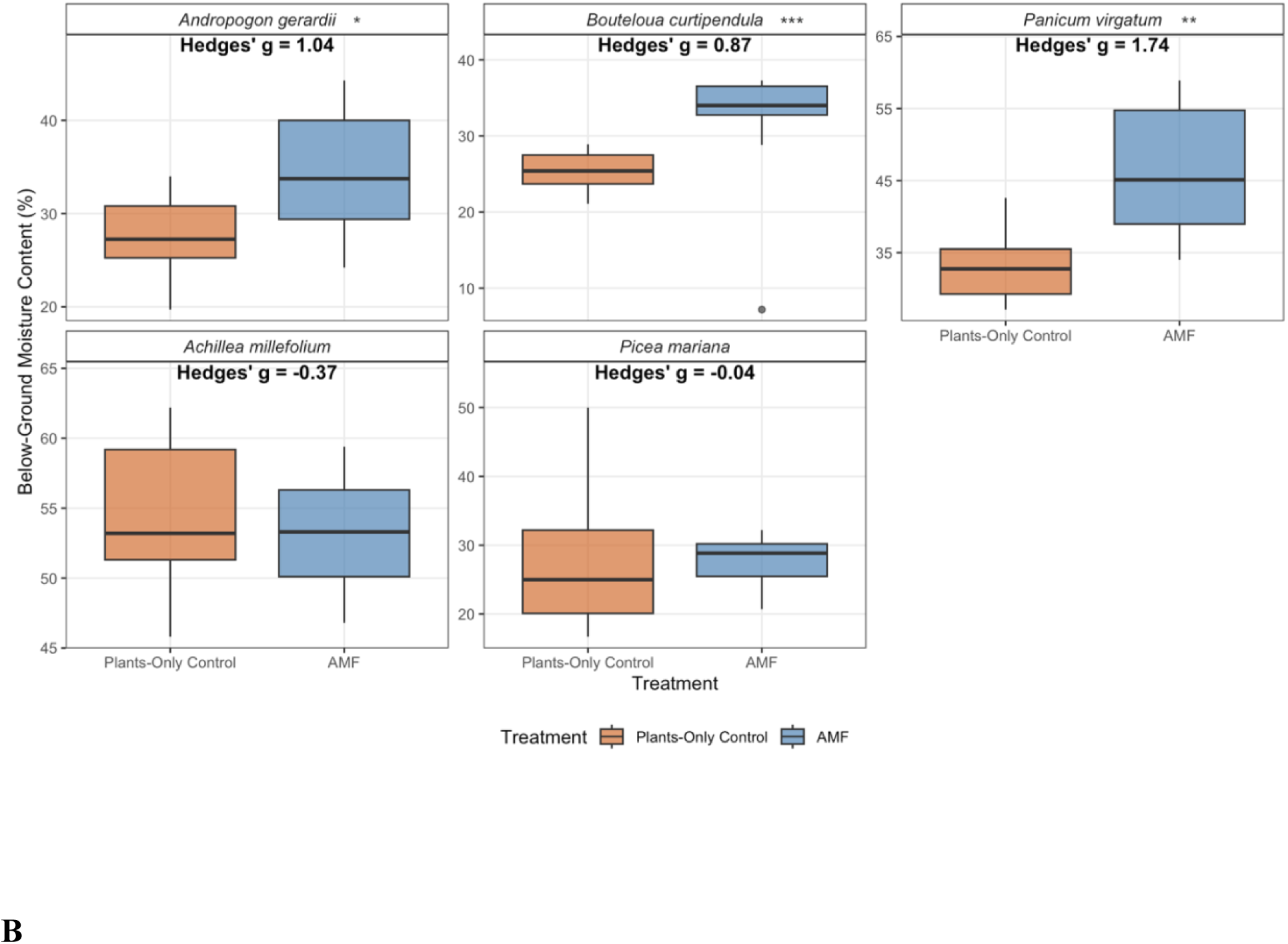
Boxplots of moisture content (%) in the five plant species under the Plants-Only Control and AMF treatments in the 25,700 mg/kg TPH soil. Mann-Whitney *U* test significant pair-wise comparisons are represented by asterisks beside plant species names in each facet (*p* < 0.001 = "***", *p* < 0.01 = "**", *p* < 0.05 = "*"). Hedge’s *g* indicate effect size comparisons between the treatments. **A.** Above-ground moisture. **B.** Below-ground moisture.

### 3.4 Plant height (January 2023 – 2025)

There was significant improvement in plant (cm) in both the <120 mg/kg and 25,700 mg/kg TPH soils under the AMF treatment (Fig. 10; both *p* < 0.05). AMF had large positive effects on plant height in both soils (*g* = 0.59 and 0.63). The median height of plants increased by 33% in the <120 mg/kg TPH soil and 54% in the 25,700 mg/kg TPH soil. In the 25,700 mg/kg TPH soil, AMF inoculation increased plant height non-significantly in all species (Mann-Whitney *U* tests: all *p* > 0.05), except *P. mariana* (Fig. S4; *p* = 0.03; *g* = 2.06).

### 3.5 Plant biomass (June 2023 – 2025)

There was a significant decrease in above-ground dry biomass (g) in the 12,600 mg/kg TPH soil compared to the <120 mg/kg TPH soil (Fig. S5; Mann-Whitney *U* test: *p* = 0.005), with a moderate effect size (Hedge’s *g* = -0.69). Below-ground dry biomass showed a non-significant decrease (*p* = 0.23) with a small effect (*g* = -0.40). Root: shoot biomass ratios were not significantly different between the two soils (*p* = 0.99; *g* = 0.03), with root growth favored over shoot growth in both soils by approximately three-fold in both the <120 mg/kg TPH (3.2-fold) and 12,600 mg/kg TPH (3.3-fold) soils.

Grass-specific above-ground biomass responses to the 12,600 mg/kg TPH soil were observed (Fig. 8). *A. gerardii* did not show statistically significant differences in dry above-ground biomass between the <120 and 12,600 mg/kg TPH soils (Mann-Whitney *U* test: *p* = 0.30). In contrast, *B. curtipendula* exhibited significant reductions (*p* = 0.002) in above-ground biomass in the 12,600 mg/kg TPH soil with a large negative effect (*g* = -1.01). *A. gerardii* showed no significant differences in below-ground biomass between the <120 and 12,600 mg/kg TPH soils (*p* = 0.68), with only a small negative effect observed with PHC contamination (g = - 0.18). *B. curtipendula* exhibited a significant (*p* = 0.003) and large reduction in below-ground biomass in the 12,600 mg/kg TPH soil (*g* = -1.02).

**Fig. 7.**
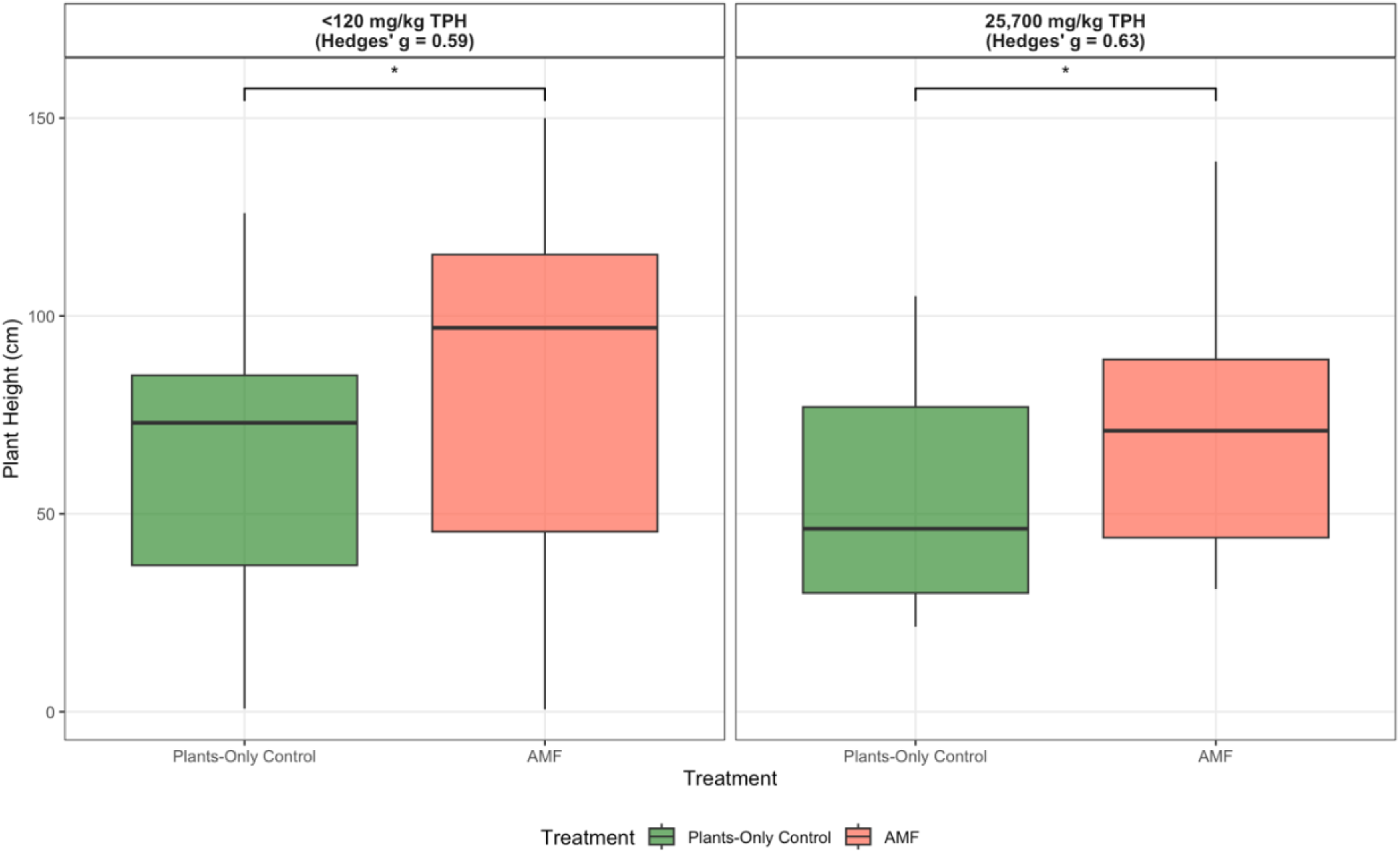
Boxplots of plant height (cm) in the <120 mg/kg TPH and 25,700 mg/kg TPH site soils under the Plants-Only Control and AMF treatments. Brackets and asterisks above boxplots indicate significant Mann-Whitney *U* test pairwise comparisons (*p* < 0.001 = "***", *p* < 0.01 = "**", *p* < 0.05 = "*"). Hedge’s *g* indicate effect size comparisons between the treatments.

**Fig. 8.**
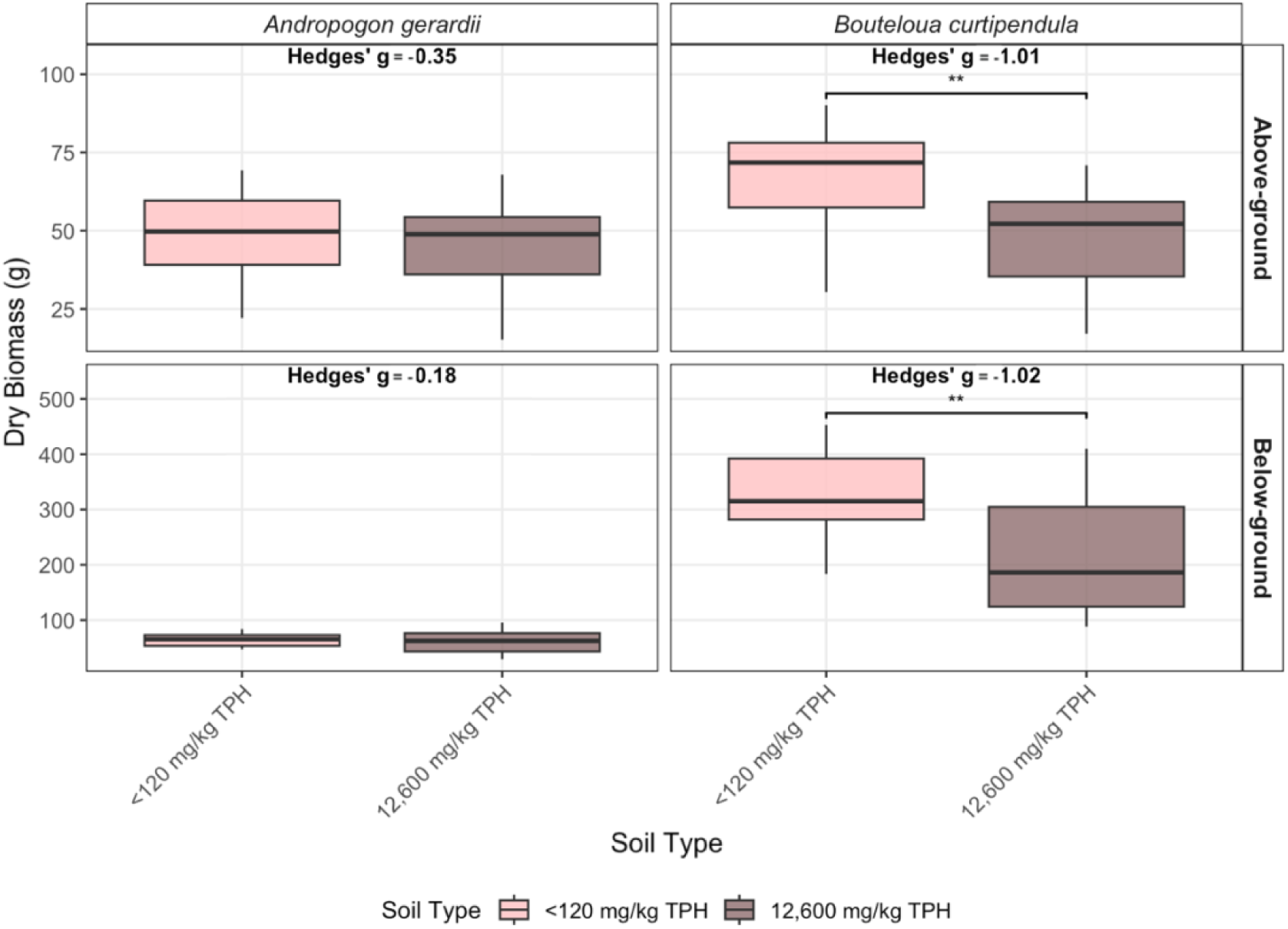
Boxplots of dry biomass (g) of *Andropogon gerardii* and *Bouteloua curtipendula* in the <120 mg/kg TPH and 12,600 mg/kg TPH soils. Boxplots are faceted by above-ground (top facets) and below-ground (bottom facets) biomass compartments and species (*A. gerardii* = left facets; *B. curtipendula* = right facets). Brackets and asterisks above boxplots indicate significant Mann-Whitney *U* test pairwise comparisons between soil type (*p* < 0.001 = "***", *p* < 0.01 = "**", *p* < 0.05 = "*"). Hedge’s *g* indicate effect size comparisons between the soils.

Kruskal-Wallis tests indicated highly significant treatment effects on above-ground biomass (g) in both soils (both *p* < 0.001), with treatment explaining 56% of the variance (η²H = 0.56) in the <120 mg/kg TPH soil and 80% of the variance (η²H = 0.80) in the 12,600 mg/kg TPH soil. Mann-Whitney *U* tests and Hedge’s *g* calculations on above-ground dry biomass (g) revealed significant treatment effects compared to the Plants-Only Control within each soil type (Fig. 9A). In the <120 mg/kg TPH soil, AMF treatment showed significant improvements over the Plants-Only Control (*p* = 0.007) with a large positive effect (*g* = 1.10), while *B. subtilis* enhanced above-ground biomass (*p* < 0.001) with a larger effect (*g* = 1.65). The combined AMF + *B. subtilis* treatment produced the most substantial improvements in above-ground biomass (*p* < 0.001; *g* = 1.91).

**Fig. 9.**
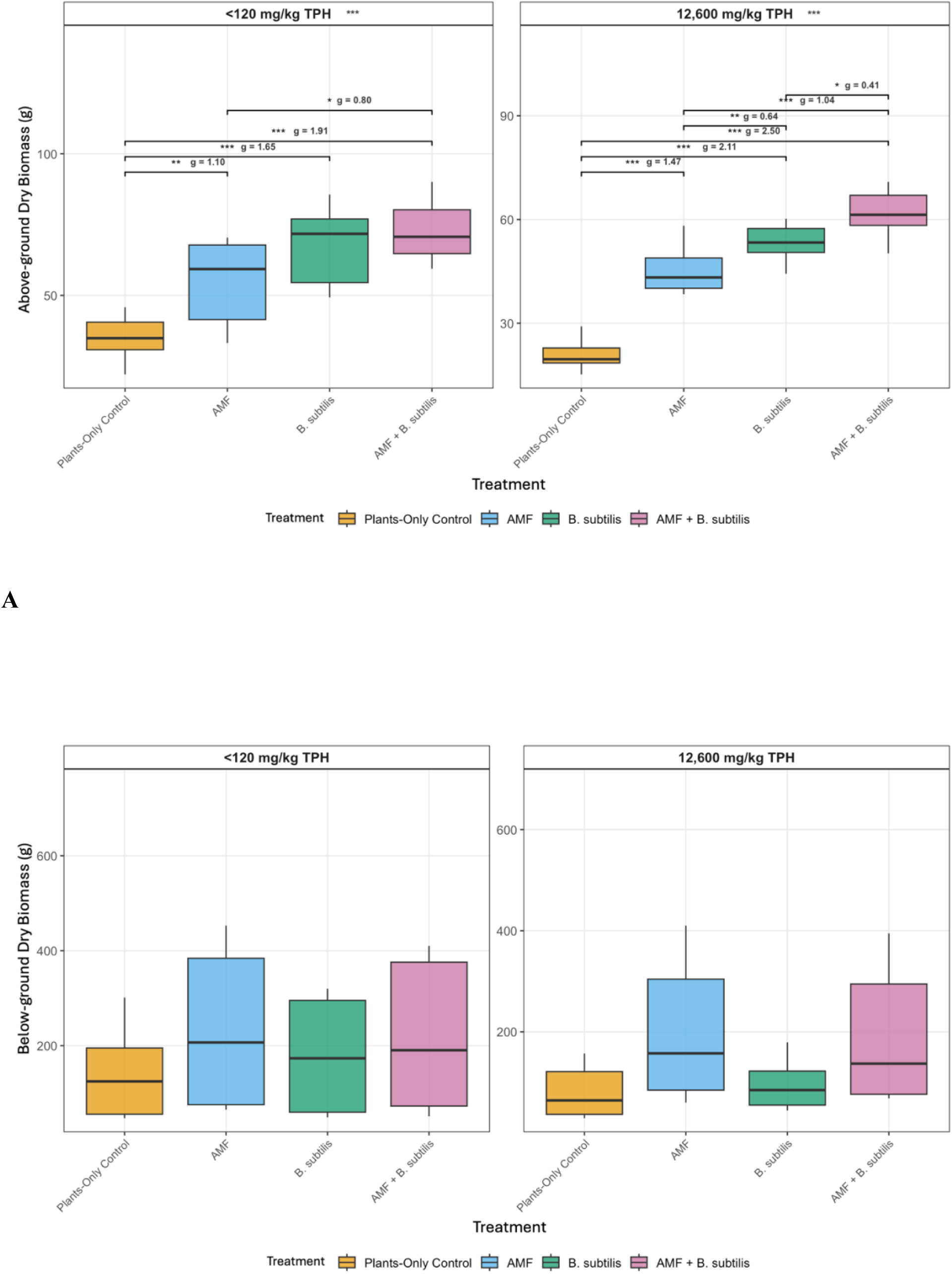
Boxplots of above- and below-ground dry biomass (g) in the Plants-Only Control, AMF, *B. subtilis* and AMF+*B. subtilis* treatments in the <120 mg/kg TPH (left facets) and 12,600 mg/kg TPH (right facets) soils. Asterisks annotated beside soil type within each facet indicate significant treatment effects (Kruskal-Wallis test: *p* < 0.001 = "***", *p* < 0.01 = "**", *p* < 0.05 = "*"). Brackets and asterisks above boxplots indicate significant Mann-Whitney *U* test pairwise comparisons between treatments. Hedge’s *g* indicate effect size comparisons between treatments. **A.** Above-ground biomass. **B.** Below-ground biomass.

**Fig. 10.**
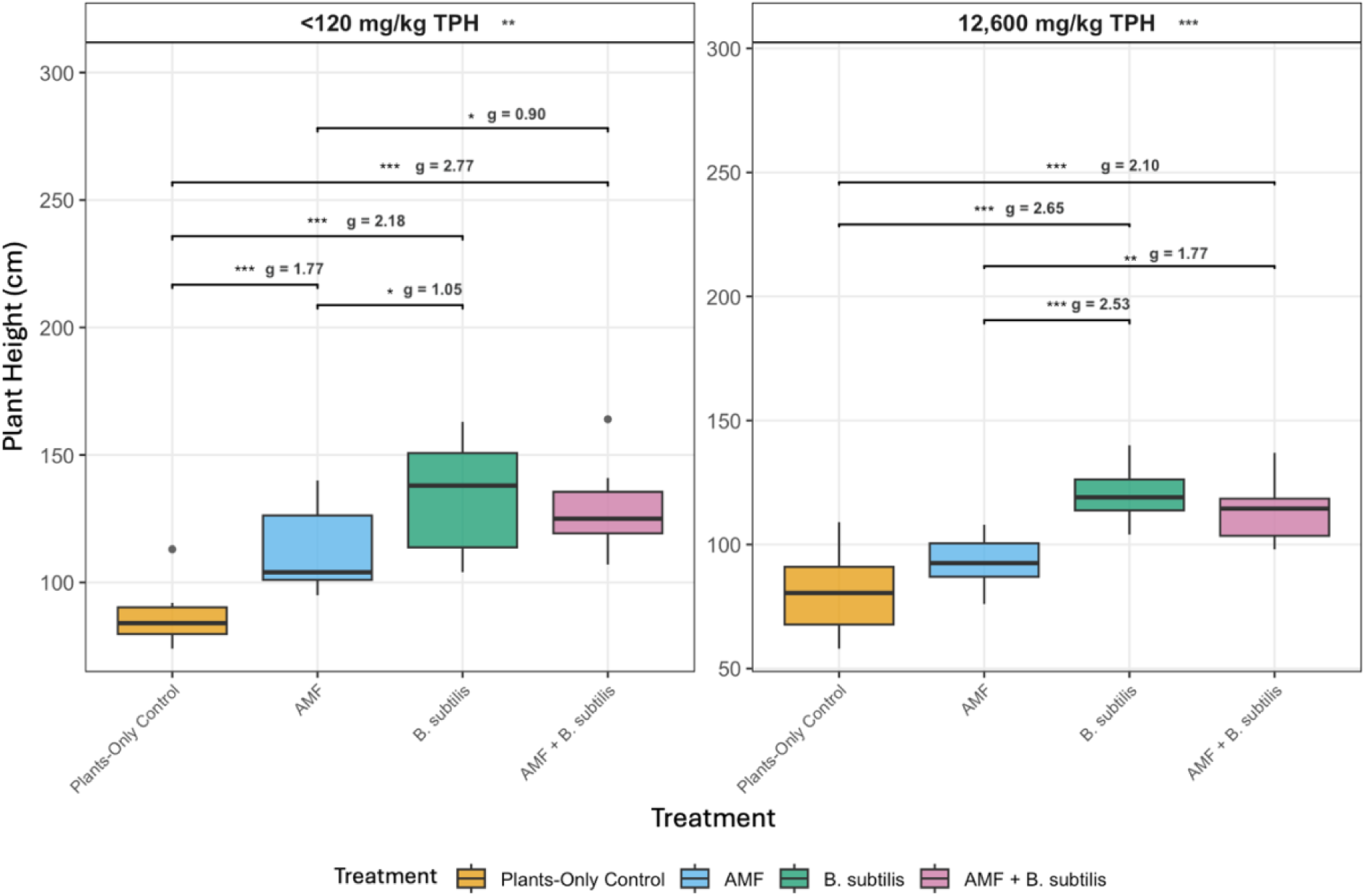
Boxplots of plant height (cm) in the <120 mg/kg TPH and 12,600 mg/kg TPH site soils under the Plants-Only Control and AMF treatments. Asterisks annotated beside soil type within each facet indicate significant treatment effects (Kruskal-Wallis test: *p* < 0.001 = "***", *p* < 0.01 = "**", *p* < 0.05 = "*"). Brackets and asterisks above boxplots indicate significant Mann-Whitney *U* test pairwise comparisons between treatments. Hedge’s *g* indicate effect size comparisons between the treatments.

Treatment effects were even more pronounced in the 12,600 mg/kg TPH soil, where AMF (*p* < 0.001; *g* = 0.64), *B. subtilis* (*p* < 0.001; *g* = 1.47), and AMF+*B. subtilis* (*p* < 0.001; *g* = 2.11) all demonstrated significant positive improvements in above-ground biomass compared to the Plants-Only Control (Fig. 9A). *B. subtilis* showed moderate improvements over AMF alone in the <120 mg/kg TPH soil (*g* = 0.80), while the AMF+*B. subtilis* treatment significantly outperformed both AMF (*g* = 1.04) and *B. subtilis* alone (*g* = 0.41) in the 12,600 mg/kg TPH soil. Overall, treatment effectiveness on above-ground dry biomass ranked as follows: AMF + *B. subtilis* > *B. subtilis* > AMF > Plants-Only Control, with above-ground biomass improvements of 1.6- to 2.1-fold observed in the <120 mg/kg TPH soils and 2.1- to 2.9-fold in 12,600 mg/kg TPH soils.

There were no significant soil treatment effects on below-ground dry biomass compared to the Plants-Only Control in both the <120 mg/kg and 12,600 mg/kg TPH soils (Fig. 9B). In the <120 mg/kg TPH soil, the AMF treatment demonstrated the most pronounced below-ground biomass enhancement compared to the Plants-Only Control, increasing median below-ground biomass by 71%. Treatment effects were more pronounced in the 12,600 mg/kg TPH soil, where the AMF+*B. subtilis* treatment demonstrated a 2.5-fold improvement. Overall, treatment effectiveness for below-ground biomass followed the pattern: AMF ≈ AMF + *B. subtilis* > *B. subtilis* > Plants-Only Control.

### 3.5 Plant height (June 2023 – 2025)

Kruskal–Wallis tests confirmed significant treatment effects on plant height (cm) in both soils (both *p* < 0.01), with treatment explaining 56% of the total variance in the <120 mg/kg TPH soil (η²H = 0.56) and 63% of the variance in the 12,600 mg/kg TPH soil (η²H = 0.63). In the <120 mg/kg TPH soil, all three soil amendment treatments significantly improved plant height compared to the Plants-Only Control (Fig. 10). The greatest enhancement was observed in the combined AMF + *B. subtilis* treatment (*p* < 0.001, *g* = 2.77), which resulted in a 49% median increase in height over the Plants-Only Control. *B. subtilis* alone produced the largest median height gain (64% increase; *p* < 0.001, *g* = 2.18), and significantly outperformed AMF alone (*p* = 0.017, *g* = 1.05); however, the difference between *B. subtilis* and AMF + *B. subtilis* was not significant (*p* = 0.50).

In the 12,600 mg/kg TPH soil, *B. subtilis* again achieved the most substantial improvement in plant height (*p* < 0.001, *g* = 2.65), raising the median height by 48% compared to the Plants-Only Control (Fig. 10). The AMF + *B. subtilis* treatment also showed significant enhancement over the control (*p* = 0.001, *g* = 2.10, 42% median increase). AMF alone did not significantly increase plant height (*p* = 0.150, *g* = 2.53), with only a 15% median gain over control pots. Overall, treatment effectiveness for plant height followed the pattern: *B. subtilis* > AMF + *B. subtilis* > AMF > Plants-Only Control in the <120 mg/kg TPH soil, and *B. subtilis* ≈ AMF + *B. subtilis* > AMF ≈ Plants-Only Control in the 12,600 mg/kg TPH soil, with *B. subtilis* consistently demonstrating the most pronounced growth-promoting effects across both soil types.

## 4.0 Discussion

### 4.1 Phytoremediation and natural attenuation

This two-year phytoremediation study demonstrated the substantial potential of plant-based remediation systems for 25+ year weathered PHC-contaminated field soil collected from the Canadian boreal ecozone. Natural attenuation (No Plants Control), plant-enhanced biodegradation (Plants-Only Control), and a combination of AMF and *B. subtilis* ATCC 21332 treatments (AMF, *B. subtilis*, and AMF + *B. subtilis*) showed a clear hierarchy of remediation efficiency (Figs. 1A-D and 2A-D). In the 25,700 mg/kg TPH soil, natural attenuation processes alone achieved significant TPH reduction, with TPH concentration declining 24% (25,700 mg/kg to 19,600 mg/kg) and 74% (12,600 mg/kg to 3,290 mg/kg) in the No Plants Controls. The superior remediation efficacy observed in the 12,600 mg/kg TPH soil compared to the 25,700 mg/kg TPH soil reflects well-established concentration-dependent relationship in PHC biodegradation that have been documented across multiple studies and has been substantiated by the US EPA (2014). Kachieng’a and Momba, (2017) found a clear inverse relationship between initial petroleum concentration and removal efficiency. Concentration-dependent inhibition occurs because higher contaminant levels can exceed microbial metabolic capacity and create toxic conditions that impair bacterial growth and enzymatic activity. Ani and Ochin, (2018) noted that while TPH degradation can be initially favored at higher concentrations due to increased substrate availability, biological half-lives increase with rising TPH concentrations, indicating slower overall remediation kinetics.

Significant degradation of PHCs in the No Plants Control pots likely resulted from stimulation of indigenous microbial metabolism of hydrocarbon compounds (bioattenuation) through enhanced aeration during initial sieving and potting, consistent watering, and photolysis through light exposure over the experimental periods (Figs. 1A-D and 2A-D; Cai et al., 2016). Volatilization of PHCs is inhibited by cold climate in the boreal ecozone, has been shown to account for as little as 2% of PHC removal from diesel, and is primarily used for benzene, toluene, ethyl benzene and xylene (BTEX) removal (Naidu et al., 2012). In this study, the 25,700 mg/kg TPH soil contained significantly higher CCME F2 and F3 levels than federal and provincial guidelines, indicating that volatilization alone was insufficient in remediating the site soil after 25+ years (Figs. 1B-C; CCME, 2008; OME, 2011; Roy et al., 2023, 2024).

Phytoremediation significantly enhanced remediation of PHCs across all CCME fractions in this study, except F3 in the 25,700 mg/kg TPH soil, aligning with findings from earlier studies that confirm the enhancement of remediation using plants compared to natural attenuation (Figs. 1A-D; Cai et al., 2016; García-Sánchez et al., 2018). The persistence of F3 hydrocarbons is consistent with the recalcitrance of higher molecular weight aliphatics and associated polycyclic aromatic hydrocarbons (PAHs), which exhibit low aqueous solubility and strong sorption to soil organic matter (CCME, 2008; Gros et al., 2014). Aging of the 25+ years weathered site soil may have led to increased incorporation of F3 compounds into soil micropores and the formation of organo-mineral complexes, further decreasing their biodegradation potential.

Despite their inherent recalcitrance, F4 hydrocarbons were significantly reduced in all phytoremediation treatments (Figs. 1A-D and 2A-D). Roy et al., (2025) identified that Proteobacteria and Actinobacteriota dominated the 12,600 mg/kg and 25,700 mg/kg TPH soils. These phyla are known to encode specialized long-chain (>C20 effective carbon number) alkane monooxygenases (*AlkB*) variants - *LadA*, and *AlmA* - which catalyze the terminal hydroxylation of long-chain alkanes, initiating their catabolism (Semenova et al., 2023). Furthermore, Actinobacteria in cold environments have demonstrated 60–70% removal of TPH and near-complete oxidation of long-chain fractions, underscoring their pivotal role in F4 degradation in boreal habitats (Vázquez Rosas Landa et al., 2023). Together, plant-mediated bioavailability enhancement and the enzymatic capabilities of indigenous rhizobacteria may have synergized to overcome the recalcitrance of F4 compounds.

### 4.2 Most Effective Phytoremediation Treatment

AMF-based treatments (AMF in the 25,700 mg/kg TPH soil and AMF+*B. subtilis* in the 12,600 mg/kg TPH soil) were identified as most effective for remediation in this study (Figs. 1A-D and 2A-D). The superior performance of AMF in enhancing PHC remediation likely reflects the multifaceted roles of AMF in soil–plant–microbe interactions. AMF colonization increases the effective root surface area through extensive extraradical hyphal networks, thereby facilitating greater contact between roots, soil-bound hydrocarbons, and hydrocarbon-degrading microbial consortia (Smith and Read, 2008; Rajtor and Piotrowska-Seget, 2016). Upon colonization, AMF divert a substantial fraction (20%) of plant photosynthate into extraradical hyphal networks, effectively extending the root’s absorptive surface and creating ‘hotspots’ of labile carbon deposition beyond the root zone (Lanfranco et al., 2016). These networks not only improves nutrient and water uptake under PHC stress, but also enhances the delivery of labile carbon (sugars and organic acids) to the rhizosphere via hyphal exudates, stimulating indigenous PHC-degrading bacteria (Zhang et al., 2024).

In the January 2023 - 2025 experiments, AMF overall reduced TPH levels in the 25,700 mg/kg TPH soil significantly more than the Plants-Only Control (Fig. 1A). In the 12,600 mg/kg TPH soil, although insignificant in magnitude, the AMF+ *B. subtilis* and AMF treatments outperformed the *B. subtilis* treatment across both grass species (Fig. 2A). These findings underscore the significant role of AMF symbiosis in driving PHC phytoremediation. Plant-AMF symbiosis induces plant systemic responses that upregulate stress-related enzymes (peroxidases, polyphenol oxidases, and glutathione *S*-transferases) and secondary metabolites (phenolics, flavonoids, and terpenoids), which can co-metabolically accelerate the breakdown of complex aliphatic and aromatic hydrocarbon fractions (Smith and Read, 2008). These compounds not only bolster plant tolerance to hydrocarbon-induced oxidative stress but also serve as co-substrates or inducers for microbial oxygenases involved in PHC rhizodegradation. Cai et al., (2016) demonstrated that AMF-colonized ryegrass exhibited two- to three-fold higher root peroxidase activity alongside increased root exudation of simple phenolics, which enhanced co-metabolic degradation of C16–C32 (C = effective carbon number) aliphatic hydrocarbons by rhizobacteria.

Roy et al., (2025) observed that microbial biodiversity and hydrocarbon-degrading bacterial communities were primarily shaped by soil PHC levels rather than treatment or plant species. This pattern otherwise suggests that indigenous rhizobacteria respond more strongly to hydrocarbon substrate availability than to AMF colonization (Jia et al., 2023). Enhanced PHC biodegradation was attributed to elevated soil carbon levels in the 12,600 and 25,700 mg/kg TPH soils rather than AMF-mediated effects (Roy et al., 2025). These observations reveal that our understanding of AMF-rhizobacteria interactions in PHC bioremediation remains incomplete, particularly regarding the mechanisms through which these partnerships influence indigenous microbial communities and hydrocarbon bioavailability in contaminated soils (Dagher et al., 2020).

### 4.3 Most effective phytoremediation plants

This study revealed that *P. mariana* was the most successful plant in remediating the 25,700 mg/kg TPH soil (Figs. 1A-D). Furthermore, both above-ground and below-ground biomass of *P. mariana* was significantly enhanced in the 25,700 mg/kg TPH soil compared to the <120 mg/kg control (Figs. 3A-B). Elevated above-ground moisture content in the 25,700 mg/kg soil (Fig. 5A) further indicates the advanced osmotic adjustment and stomatal regulation capacity of this species, which preserve turgor and mitigate hydrocarbon-driven oxidative damage (Khan et al., 2013). In an earlier study, *P. mariana* showed the least phytotoxic response to 11,900 mg/kg TPH soil from the same Canadian boreal site examined in this study (Roy et al., 2023).

*P. mariana* synthesizes high levels of antioxidative enzymes—such as superoxide dismutase and catalase—that detoxify reactive oxygen species generated during hydrocarbon exposure, protecting membrane integrity and allowing continued biomass accumulation (García-Pérez et al., 2012). As a long-lived conifer, *P. mariana* possesses constitutively high levels of the CYP720B family of cytochrome P450 monooxygenases, enzymes that catalyze hydrocarbon-oxidation reactions and serve as precursors for diterpene resin acid biosynthesis (Bathe and Tissier, 2019). This oxidative capacity may facilitate co-metabolic transformation of PHCs into less toxic metabolites, thereby protecting cellular membranes and sustaining growth under PHC stress (Roy et al., 2023). Together, these traits likely enabled *P. mariana* to successfully grow in the 25+ years weathered PHC-contaminated soil, resulting in greater above- and below-ground biomass and enhanced moisture retention compared to other plants grown in the <120 mg/kg TPH control soil (Figs. 3A-B).

In the 12,600 mg/kg TPH soil, both *A. gerardii* and *B. curtipendula* significantly reduced PHC levels to below federal CCME and provincial thresholds across all treatments (Figs. 2A-D). The roots of both species – fibrous lateral (*A. gerardii*) and stoloniferous (*B. curtipendula*) - reached the full length of the pots (45.6 cm) across all treatments. Moderate hydrocarbon contamination stimulates root proliferation of resilient graminoids, enhancing soil microbial activity in the rhizosphere via root exudates (Merkl et al., 2005). Merkl et al., (2005) observed increased root surface area and diameter in graminoids under 2% crude oil contamination. These findings can be attributed to cortical cell radial expansion, root exoderm thickening, and/or cortical parenchyma expansion to enhance water and nutrient uptake under abiotic stress (Potocka and Szymanowska-Pułka, 2018). Increased root area provides more habitat for PHC-degrading bacteria and labile carbon sources that facilitate indigenous microbes to co-metabolize recalcitrant and bioavailable PHCs (Balasubramaniyam, 2015).

There was no significant difference in above- and below-ground biomass of *A. gerardii* in the PHC-contaminated (12,600 and 25,700 mg/kg TPH) and background soils in this study, whereas *B. curtipendula* biomass was significantly inhibited by PHC contamination (Figs. 3A-B and 8). *A. gerardii* develops a deep, robust root system with short rhizomes and extensive lateral roots that extend deeply into the soil profile (150–240 cm in field soils), facilitating plant structural stability and soil nutrient uptake during PHC stress (USDA, 2015). In contrast, *B. curtipendula* forms a dense and fibrous network with narrow scaly rhizomes and occasional stolons that favour root exploration in non-contaminated soils. As demonstrated by the increased below-ground moisture content in *B. curtipendula* growing in the 25,700 mg/kg TPH (Fig. 5B), the relatively narrow root structure of this species in soil makes it more susceptible to abiotic stressors, including waterlogging from PHC-inhibited hydraulic conductivity, than *A. gerardii* (Wernerehl and Givnish, 2005). Furthermore, *Andropogon* species possess and rapidly induce a key antioxidant enzyme involved in PHC detoxification, glutathione *S*-transferase (Ezaki et al., 2008). These mechanisms may afford *A. gerardii* a more effective first line of defense against oxidative stress, preserving membrane integrity and metabolic function under PHC contamination.

Due to the fast-growing nature of *A. gerardii*, it is recommended over *P. mariana* for rapid revegetation of PHC-contaminated sites. In field sites, *A. gerardii* reaches full height (3 metres) within three growing seasons whereas *P. mariana* can take 10-20 years to produce cones and 50+ years to achieve full growth (Black and Bliss, 1980; USDA, 2015). In this study, *A. gerardii* exhibited a height range of 58–140 cm in the 12,600 mg/kg TPH soil, 43–140 cm in the 25,700 mg/kg TPH soil, while *P. mariana* height ranged from 21-41 cm in the 25,700 mg/kg TPH soil, indicating more rapid vertical growth of *A. gerardii* within the 24-month experimental period (Figs. 10 and S4). Additionally, *A. gerardii* exhibited significantly enhanced below-ground biomass than *P. mariana* in the 25,700 mg/kg TPH soil, across both treatments (Fig. 3B; Mann-Whitney *U* tests: *p* < 0.001), The more rapid root development of *A. gerardii* may facilitate PHC remediation within deeper soil horizons within a limited remediation time period (Thomas et al., 2017). However, fully-grown indigenous *P. mariana* trees at boreal field sites may present a promising opportunity for site remediation and reclamation.

### 4.4 Overall effects on plant health

Plant species grown in PHC-contaminated soils (12,600 mg/kg and 25,700 mg/kg TPH) exhibited significant declines in both above- and below-ground biomass relative to background (<120 mg/kg TPH) soils (Figs 3A-B and S1), and a significantly higher root: shoot moisture ratio, consistent with the phytotoxic effects of hydrocarbons on vascular plants reported in the literature (Khan et al., 2013; Cai et al., 2016). The high root: shoot biomass and soil moisture ratios (>1) in all three soils examined in this study may be reflective of both plant stress resulting from both boreal soil conditions and PHC phytotoxicity (Robertson et al., 2007). Low soil carbon content was observed in the <120 mg/kg TPH soil whereas nitrogen and phosphorous deficiency was noted in the 25,700 mg/kg TPH soil (Roy et al., 2025). Under stress conditions, plants typically exhibit increased biomass allocation to roots as an adaptive strategy to optimize resource acquisition, with reduced nutrient availability and abiotic stressors consistently promoting greater root investment relative to shoot growth to enhance water and nutrient uptake capacity (Desalme et al., 2011).

Above-ground moisture content was significantly reduced all plant species, except *P. mariana* (Fig. 5A). Inoculation with AMF mitigated PHC stress, significantly enhancing plant height (Fig. 7), above and below-ground biomasses (Fig. 4), and both above- and below-ground moisture content relative to the Plants-Only Control in the 25,700 mg/kg TPH soil (Figs. 6A-B; S3). These findings highlight the capacity of AMF to improve plant nutrient acquisition and alleviate PHC inhibition of root development through improved soil nutrient uptake and soil structure stabilization (Rajtor and Piotrowska-Seget,^,^ 2016).

*B. subtilis* inoculation produced significant enhancements in above- and below-ground biomass compared to the Plants-Only Control and AMF treatments in 12,600 mg/kg TPH soil (Fig. 9A). Furthermore, this treatment resulted in the highest median plant growth across treatments (Fig. 10). This response aligns with known PGPR mechanisms—phytohormone production, ACC deaminase activity, and enhanced phosphorous solubilization in soil—which collectively alleviate stress-induced ethylene accumulation in plants and improve plant nutrient availability under PHC toxicity (Tsotetsi et al., 2022). These mechanisms are particularly augmented in PHC-contaminated boreal soils, where hydrocarbon toxicity often coincides with nutrient limitations that exacerbate plant stress (Gamalero and Glick, 2015).

Roy et al., (2025) observed that *B. subtilis* was not of the 30 most abundant genera found in the 12,600 mg/kg TPH soils after one year of inoculation. Furthermore, this treatment did not enhance PHC remediation compared to the Plants-Only Control in this study (Figs. 2A-D), suggesting that *B. subtilis* may have been outcompeted by indigenous rhizobacteria or taken into the plant roots as an endophyte. The abundance of indigenous PHC-degrading bacteria predatory phyla, Myxococotta, in the soil suggests outcompetition through niche competition and predation (Müller et al., 2014; Roy et al., 2025). However, the significant improvement in plant above-ground biomass in the 12,600 mg/kg TPH soil under the *B. subtilis* treatment but no observed improvement in below-ground biomass may be suggestive of endophyte colonization - although the strain of *B. subtilis* used in this study (ATCC 21332) is not well-documented for its endophytic abilities (Figs. 9A-B; Bolivar-Anillo et al., 2021).

Overall, the *B. subtilis* treatment did not enhance PHC remediation relative to the AMF or AMF+*B. subtilis* treatments (Figs. 2A-D), but it did significantly improved plant above-ground biomass and plant height in the 12,600 mg/kg TPH soil (Figs. 9A and 10). The *B. subtilis* treatment is recommended for improving plant health during phytoremediation while the AMF treatment is recommended for enhanced phytoremediation of PHCs during field trials. Given these complementary functions, and effectiveness of the AMF+*B. subtilis* treatment in remediating the 12,600 mg/kg TPH soil (Fig. 2A), a tiered approach may be used. Specifically, *B. subtilis* inoculation may be utilized for early plant establishment from seed in a greenhouse while AMF inoculation is amended into PHC-contaminated field site *in situ* before planting. Field experiments using both AMF and *B. subtilis* also merits exploration, as co-inoculation using *B. subtilis* and AMF has improved crop yield in field trials (Nanjundappa et al., 2019).

## 5.0 Conclusion

This two-year greenhouse study demonstrated that phytoremediation of 25+ years weathered PHC-contaminated field soil (25,700 mg/kg and 12,600 TPH) can be significantly enhanced by the strategic use of plant–microbe partnerships. While natural attenuation alone achieved significant hydrocarbon reduction, planting indigenous and naturalized boreal plant species accelerated remediation and brought the soils containing 12,600 mg/kg TPH to below both federal and provincial regulatory thresholds. Among the four treatments assessed, inoculation with AMF-based treatments yielded the greatest TPH removal: AMF in the 25,700 mg/kg TPH soil (74% mean removal) and AMF+*B. subtilis* in the 12,600 mg/kg TPH soil (96%). *Picea mariana* emerged as the most effective species for remediation of the 25,700 mg/kg TPH soil while tufted grass species (*Andropogon gerardii* and *Bouteloua curtipendula*) remediated the 12,600 mg/kg TPH soils to federal and provincial guideline levels within 17 months. *A. gerardii* above- and below-ground dry biomass was not impaired by PHC contamination, making it an ideal candidate for rapid revegetation and effective phytoremediation of PHC-contaminated soils. The PGPR, *B. subtilis* ATCC 21332, significantly promoted plant growth through enhanced above-ground biomass and plant height. Combining *B. subtilis* for robust seedling establishment with AMF for field-scale remediation offers a promising strategy. These findings support the advancement of field-scale phytoremediation trials at PHC-contaminated sites across the boreal ecozone, where AMF- and *B. subtilis*-enhanced phytoremediation could provide cost-effective and environmentally sustainable alternatives to conventional excavation and disposal methods. Future research should prioritize long-term field trials that monitor both hydrocarbon degradation rates and ecosystem recovery indicators, establishing standardized protocols for healthy plant establishment to facilitate broader implementation across PHC-contaminated sites in boreal ecosystems.

## Acknowledgements

We thank Officer Courtney Drew, Officer Victoria Huppe, and Officer Cadet Hunter Patterson for their assistance in harvesting the greenhouse plants and taking biomass measurements. We also members of the Analytical Service Unit at Queen’s University, including Dr. Graham Cairns, Paula Whitley, and Mesha Thompson, for analytical assistance. We also thank Linda Eastcott for her support in conducting this research.

