## Supplemental Figures S1 to S4 for "Phytoremediation of 25+ Years Weathered Petroleum Contamination with Symbiotic Arbuscular Mycorrhizal Fungi and *Bacillus subtilis* ATCC 21332"

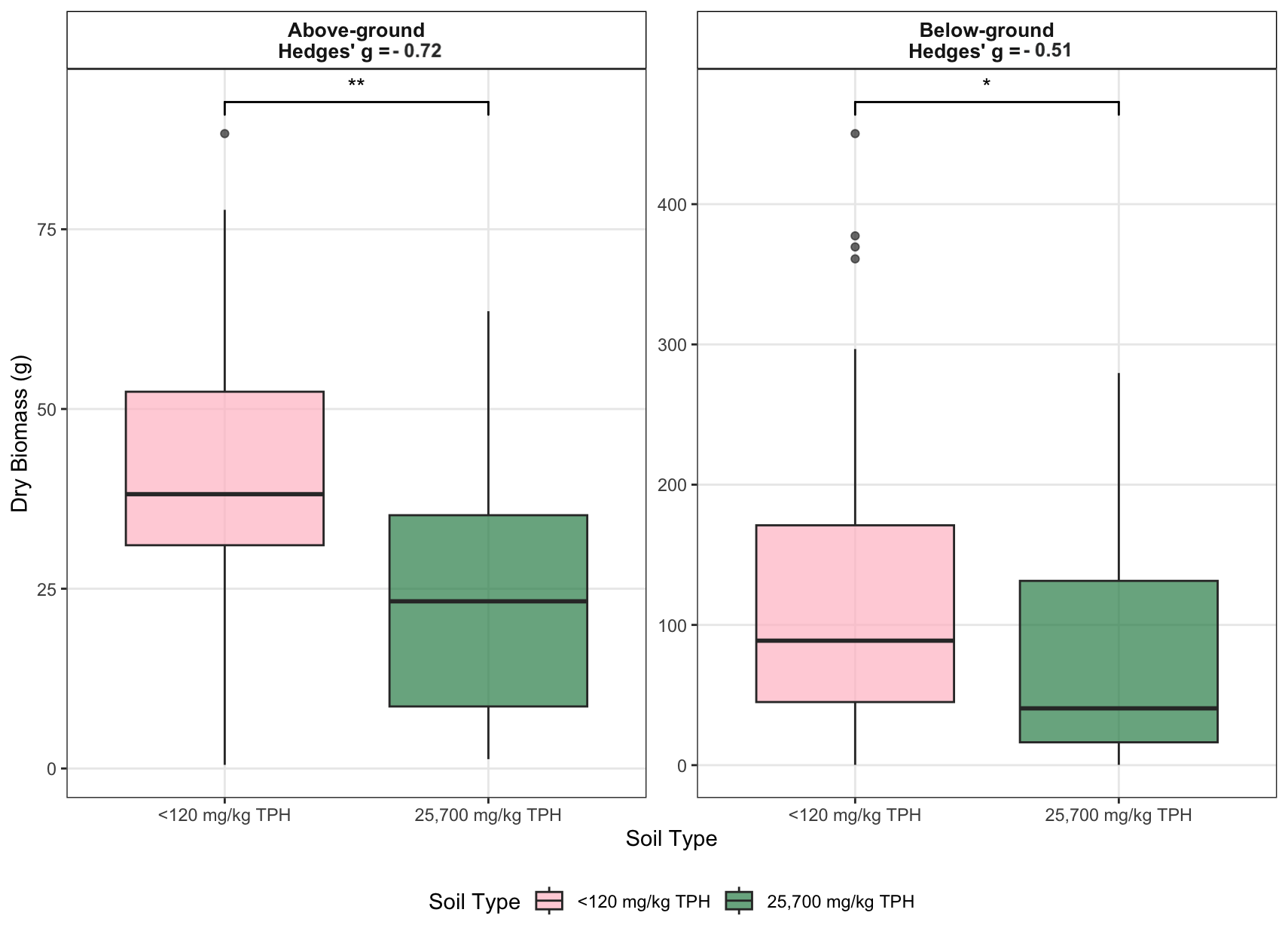

**Fig. S1.** Boxplots of dry above- and below-ground biomasses (g) in the <120 mg/kg TPH and 25,700 mg/kg TPH site soils following two-years of phytoremediation. Brackets and asterisks above boxplots indicate significant Mann-Whitney *U* test pairwise comparisons between the soils (*p* < 0.001 = "***", *p* < 0.01 = "**", *p* < 0.05 = "*"). Hedge’s *g* indicate effect size comparisons between the site soils.

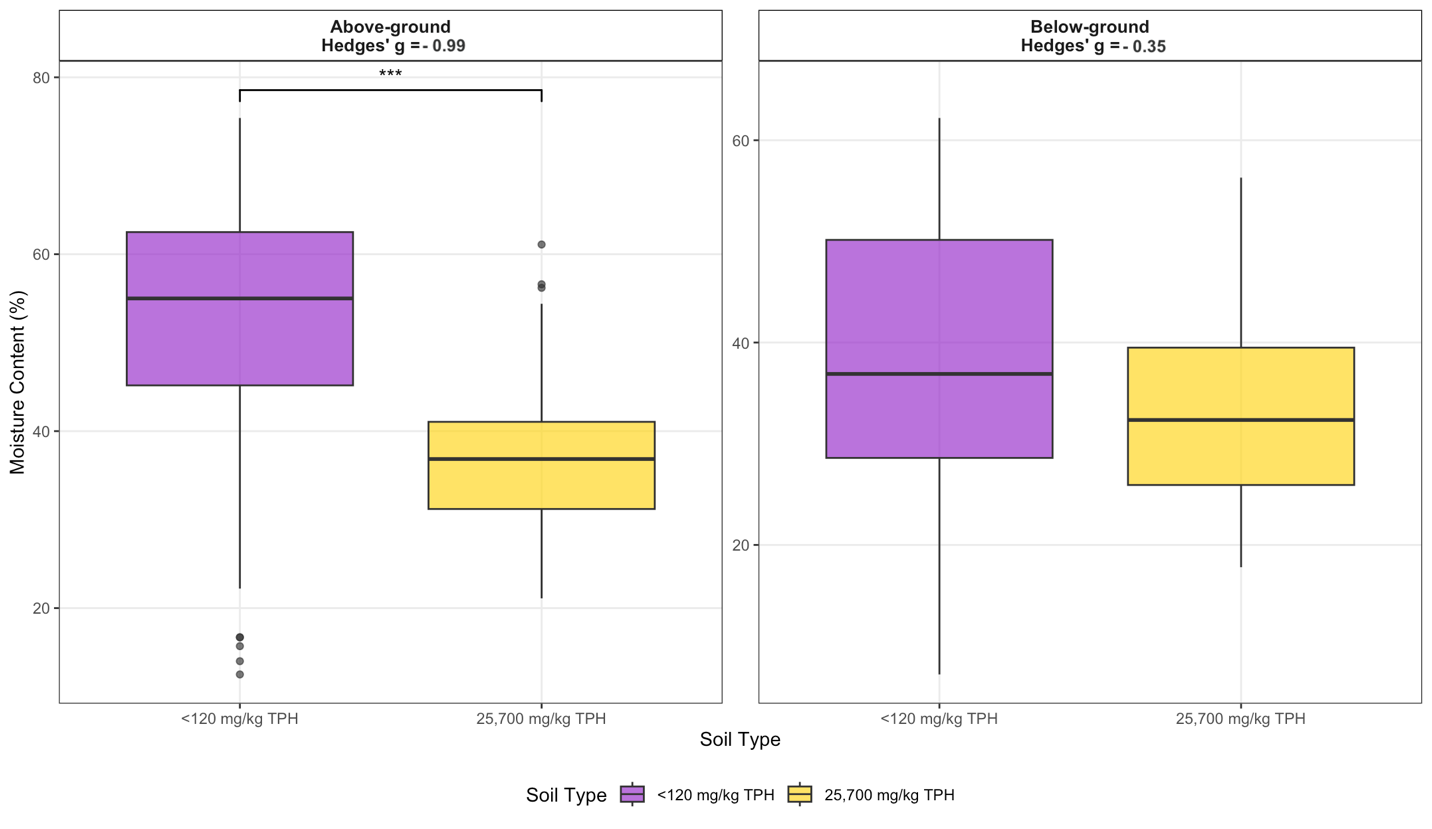

**Fig. S2.** Boxplots of above- and below-ground plant moisture content (%) in the <120 mg/kg TPH and 25,700 mg/kg TPH site soils. Brackets and asterisks above boxplots indicate significant Mann-Whitney *U* test pairwise comparisons (*p* < 0.001 = "***", *p* < 0.01 = "**", *p* < 0.05 = "*"). Hedge’s *g* indicate effect size comparisons between the site soils.

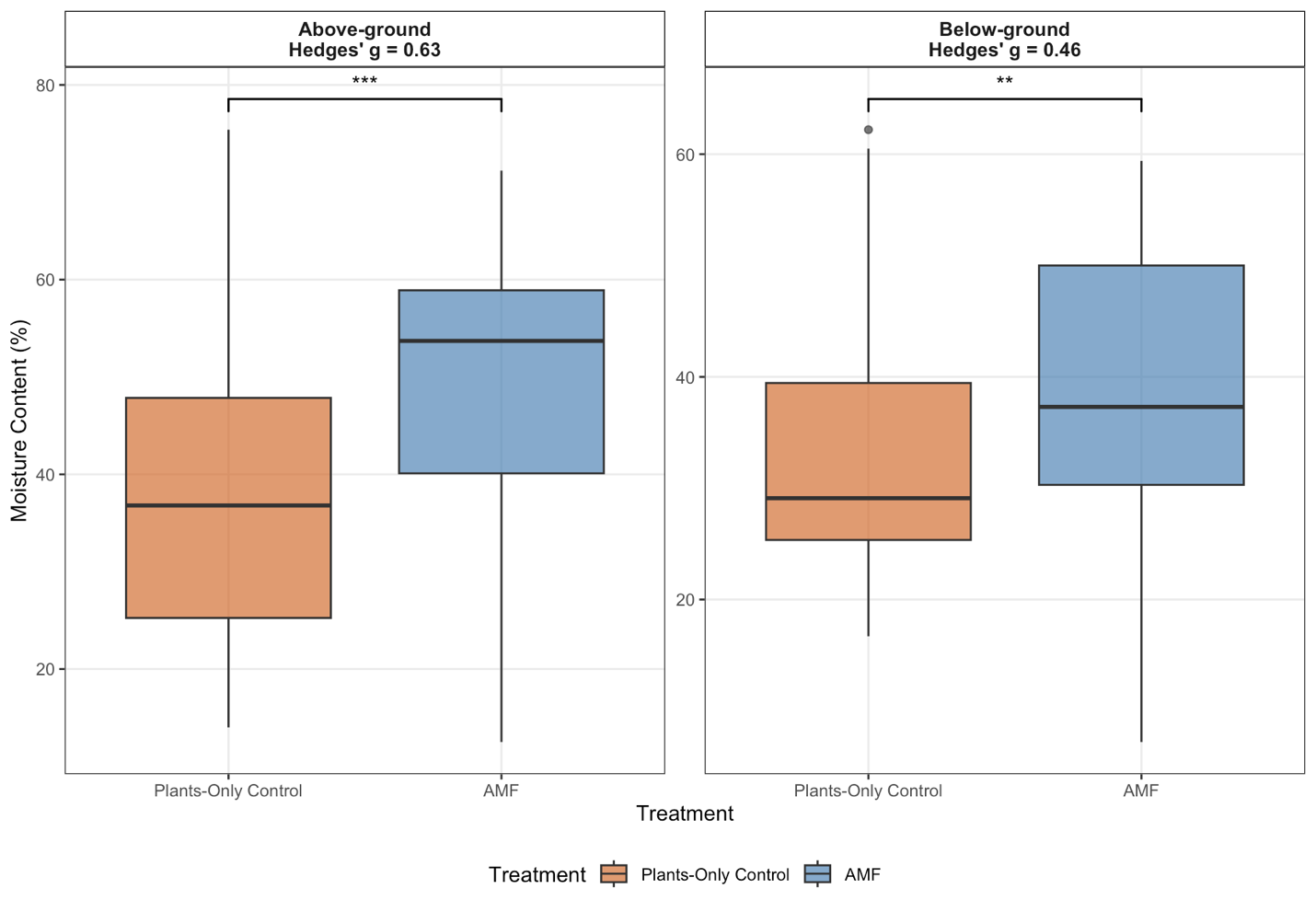

**Fig. S3**. Boxplots of above- and below-ground plant moisture content (%) in the Plants-Only Control and AMF treatments (25,700 mg/kg TPH soil). Brackets and asterisks above boxplots indicate significant Mann-Whitney *U* test pairwise comparisons (*p* < 0.001 = "***", *p* < 0.01 = "**", *p* < 0.05 = "*"). Hedge’s *g* indicate effect size comparisons between the treatments.

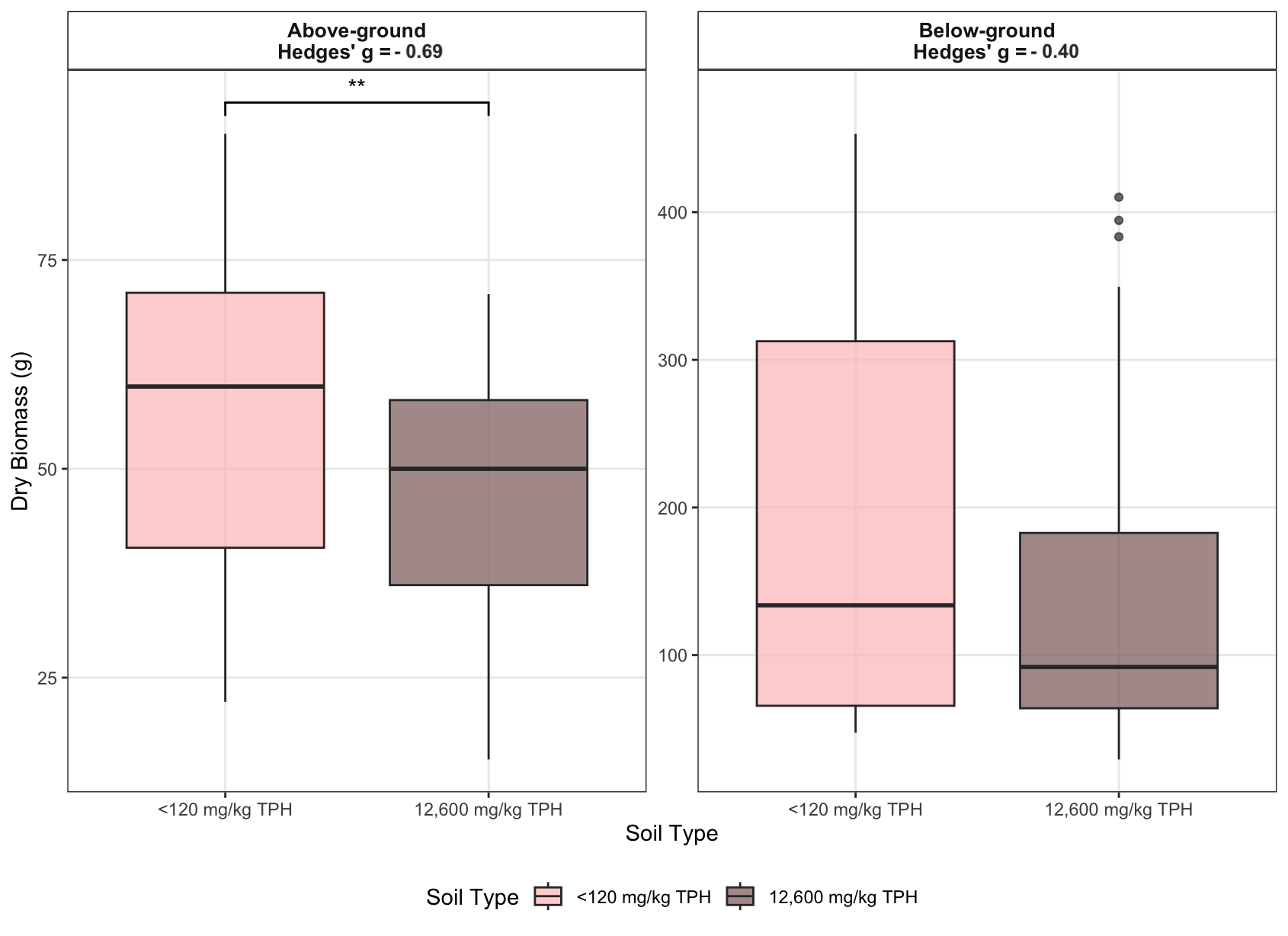

**Fig. S4.** Boxplots of dry above- and below-ground biomasses (g) of *Andropogon gerardii* and *Bouteloua curtipendula* in the <120 mg/kg TPH and 12,600 mg/kg TPH soils following two-years of phytoremediation. Brackets and asterisks above boxplots indicate significant Mann-Whitney *U* test pairwise comparisons between the soils (*p* < 0.001 = "***", *p* < 0.01 = "**", *p* < 0.05 = "*"). Hedge’s *g* indicate effect size comparisons between the soils.
